# Evolutionary rescue during extreme drought

**DOI:** 10.1101/2024.10.24.619808

**Authors:** Daniel N. Anstett, Julia Anstett, Seema N. Sheth, Dylan R. Moxley, Haley A. Branch, Mojtaba Jahani, Kaichi Huang, Marco Todesco, Rebecca Jordan, Jose Miguel Lazaro-Guevara, Loren H. Rieseberg, Amy L. Angert

## Abstract

Populations declining due to climate change may need to evolve to persist. While evolutionary rescue has been demonstrated in theory and the lab, its relevance to natural populations facing climate change remains unknown. Here we link rapid evolution and population dynamics in scarlet monkeyflower, *Mimulus cardinalis*, during an exceptional drought. We leverage whole-genome sequencing across 55 populations to identify climate-associated loci. Simultaneously we track demography and allele frequency change throughout the drought. We establish range-wide population decline during the drought, geographically variable rapid evolution, and variable population recovery that is predictable by standing genetic variation in and rapid evolution at climate-associated loci. These findings demonstrate evolutionary rescue in the wild, showing that genetic variation at adaptive, but not neutral loci, predicts population recovery.

## Main Text

Climate change is leading to population declines due to increasing severity and duration of extreme events (*1-3*). Reversing such declines may depend on evolutionary rescue, where evolution leads to demographic recovery (*4*). Evolutionary rescue has extensive theoretical support (*5, 6*) and elegant lab demonstrations in microorganisms (*7-9*), but evolutionary rescue has not been directly demonstrated in wild populations in response to climate change. The best evidence for evolutionary rescue outside the lab is herbicide resistance in response to modern agriculture (*10, 11*), transmissible cancer resistance in Tasmanian devils (*12*), and adaptation to pollution in killifish (*13*). However, these studies have not provided direct evidence of the link between allele frequency change and population demography. Documenting evolutionary rescue in the wild requires demonstrating (1) demographic decline driven by an environmental perturbation, (2) rapid adaptation during population decline, and (3) demographic recovery that results from evolutionary change (Fig. 1A).

**Fig 1.**
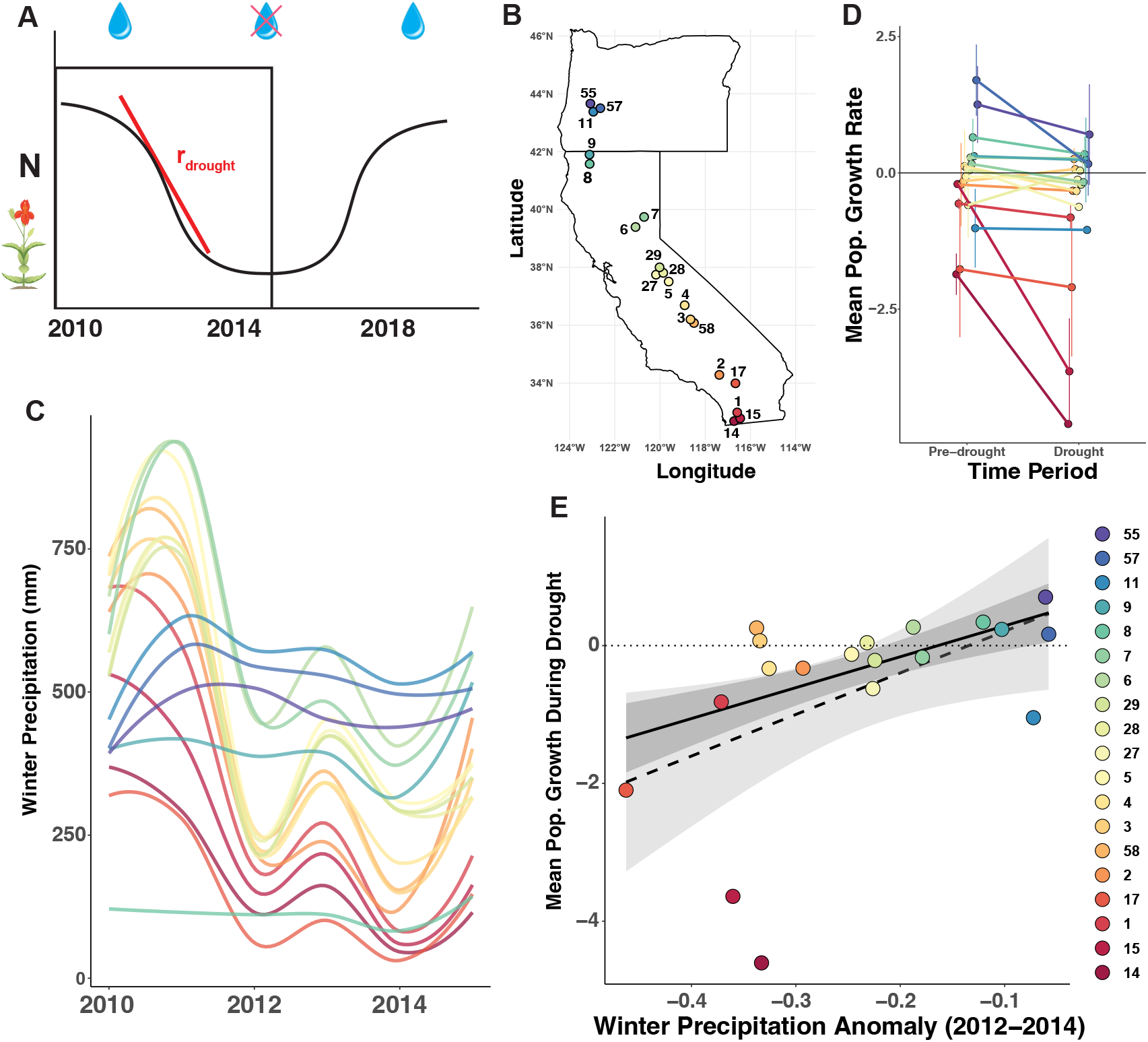
Drought-induced demographic decline. (**A**) Decline phase of evolutionary rescue, indicated by the rectangle. r_drought_ represents mean growth rate (*r*) during the drought time period (2012-13, 2013-14, 2014-15). (**B**) Map of 19 *M. cardinalis* populations considered during demographic surveys. Rainbow colouring indicates latitudinal position (red colors = lower latitudes, blue colors = higher latitudes). Population codes are given for each population (Table S3). (**C**) Mean annual precipitation (mm) across 19 populations of *M. cardinalis* before and during the record-setting drought. Each curve represents the precipitation time series for one population. (**D**) Mean *r* during pre-drought (2010-11, 2011-12), and drought time periods shown as latitudinally colored points with lines connecting the two time periods for a given population. The horizontal black line depicts rate of replacement (*r* =0); values below this line indicate a population that is projected to decline. Points are jittered to minimize overlap. (**E**) During severe drought, populations with greater winter precipitation anomaly (more negative values) experienced lower mean intrinsic growth rates (P=0.02, R^2^=0.22). Each point is one of the 19 populations where demography was assessed colored again by latitude. Solid lines and darker shading depict predicted the slope and its 95% confidence intervals from robust regression. Dashed line and lighter shading depict the slope and its 95% confidence intervals from ordinary least squares regression. The dotted horizontal line depicts rate of replacement (*r* =0).

We demonstrate evolutionary rescue during an exceptional drought in a riparian plant, scarlet monkeyflower (*Mimulus cardinalis*), by leveraging extensive demographic and landscape genomics datasets. *Mimulus cardinalis* is a perennial herb present in lowland and mountainous habitats from northern Baja California, Mexico through southern Oregon, USA (Fig. 1B). Much of the species’ range has been impacted by a 22-year megadrought (*14*), with the 2012-2015 drought representing the most extreme four-year drought in more than 10,000 years in California (*15*) (Fig. 1C). The frequency of these “exceptional” drought events is projected to rise with climate change (*2, 16*) increasing the vulnerability of California and many other Mediterranean-type ecosystems to drought stress. Because *M. cardinalis* occurs along an aridity gradient, and shows evidence of differential physiological and life history adaptation across this gradient (*17, 18*), the species is an ideal system to study geographical variation in drought adaptation over space and time.

We first assessed the impact of extreme drought on *M. cardinalis* population dynamics. Through annual field surveys from 2010 to 2018, we collected demographic data on survival, growth, reproduction, and recruitment from 19 populations along a North-South transect encompassing most of the range of *M. cardinalis* (Fig. 1B). We used these data to infer population growth rate (*r*) through integral projection modeling (IPM) (Supplementary Methods) (*19*). Eleven populations were projected to be in decline (*r*<0) during the extreme drought (Fig. 1D; Fig. S1A, C, Table S1). Despite considerable heterogeneity in population growth rates created by idiosyncratic flood and fire disturbances, mean *r* was significantly lower during drought than during the two years prior (Wilcoxon paired rank-sum test, V=151, P=0.003). Greater anomalies in winter precipitation were a strong predictor of lower mean population growth rate during the drought (Fig. 1E; Table S2), further implicating drought stress as a likely cause of these population declines.

Having established demographic decline during the exceptional drought, we then leveraged an extensive spatiotemporal collection of seed and leaf tissue to (a) identify single nucleotide polymorphisms (SNPs) associated with historical climate and (b) track their evolution during the contemporary drought (Fig. 2). First, we identified SNPs associated with 30-year pre-drought climate averages (1980-2009) by sequencing a “baseline” dataset of 55 populations (Fig. 2B). These populations were sampled across the entire range of *M. cardinalis* from 2007 to 2011, a baseline time period just prior to the exceptional 2012-2015 drought (Fig. 1C, Fig. S2 Table S3). We hypothesized that <u>historical selection varied spatially according to consistent differences in historical precipitation and temperature (Fig. 1C), and predicted a greater frequency but lower segregating diversity of drought-adapted alleles in the south.</u> The baseline dataset is composed of whole-genome sequences of 347 individuals at 6 to 21X coverage, identifying a total of 2,156,443 SNPs. We combined SNP-based (*20*) and window-based (*21*) Genotype-Environment Associations (GEAs) to identify SNPs with strong association with the historical climate (Fig. 2C). To perform these analyses, we used 9 climate variables (*22*) that quantify annual and seasonal differences in temperature, precipitation and moisture deficit across the species range. This identified a total of 605 unique climate-associated SNPs from across the genome (Fig. 2D; Fig.S3-4). Unique SNPs were further thinned to account for linkage (*23*) and monotonic relationships with climate yielding 215 SNPs, ranging from 18 to 54 SNPs per climate variable (Table S4). Populations entered the drought with different allele frequencies and genetic diversity at these loci, with southern populations harboring less standing variation at climate-associated loci despite having higher genome-wide variation compared to northern populations (Fig. S5).

**Fig 2.**
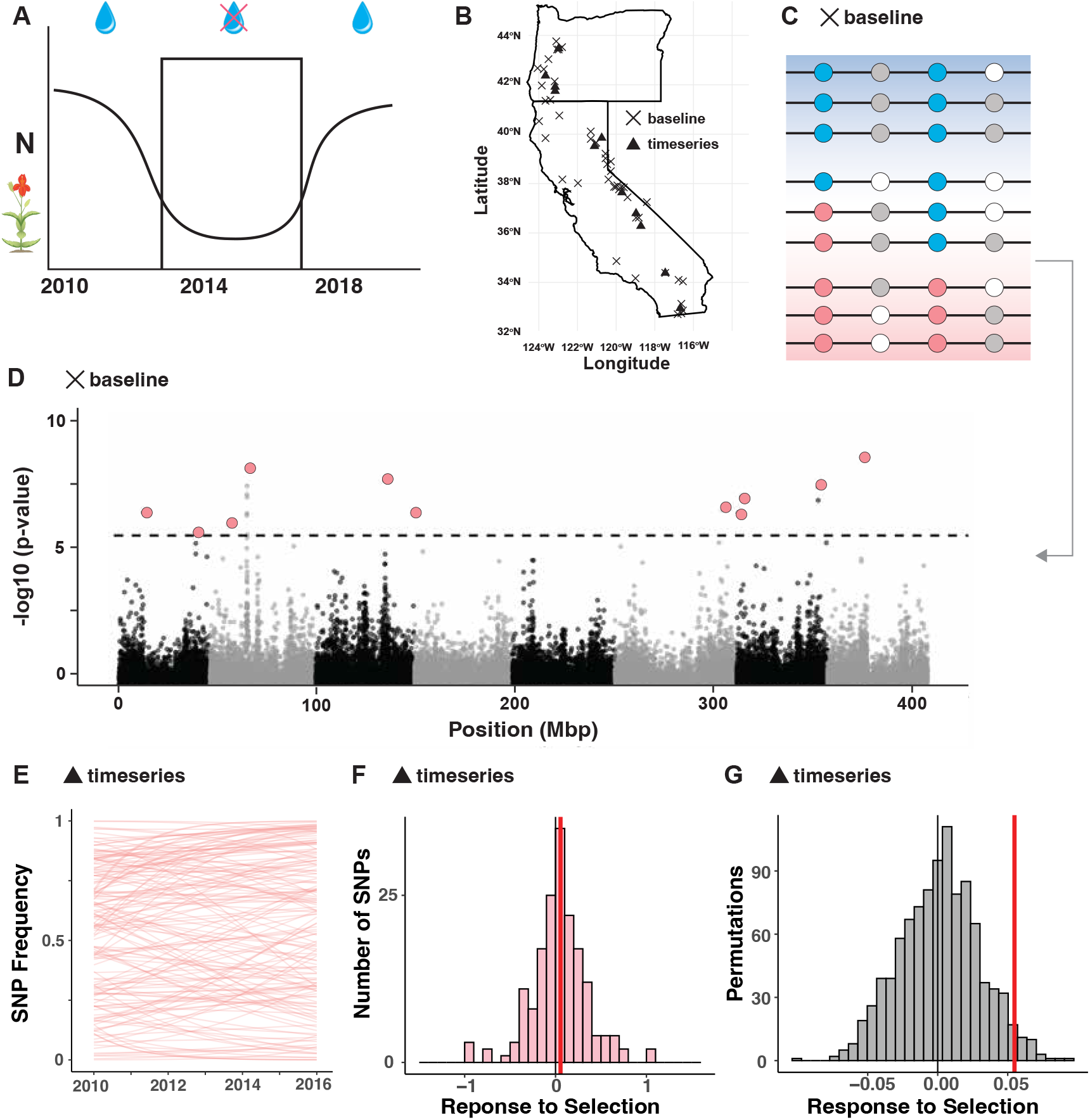
Rapid adaptation from standing genetic variation. Genotype-Environment Association (GEA) identifies Single Nucleotide Polymorphisms (SNPs) associated with spatial variation in historical climate, whose frequencies are then tracked over time during the (A) rapid evolution phase of evolutionary rescue, indicated by the rectangle. (B) Fifty-five “Baseline” *M. cardinalis* populations (all symbols, X are baseline only) were sampled prior to record-setting drought (2007-2011) for GEA. Eleven “Time Series” populations (triangle symbols) were further sampled annually from 2010-2016, encompassing the extreme drought. (C) GEA on Baseline samples identifies SNPs associated with historical (1980-2009) spatial variation in temperature and precipitation. Cartoon depicts a climate gradient (cool and wet in blue to warm and dry in pink) with populations (horizontal lines) showing differences in the most prevalent SNP at four loci. Two climate-associated loci (pink and blue alleles) and two non-climate associated loci (grey and white alleles) are shown. In this cartoon, SNP associated with warmer and drier conditions (pink alleles) are more prevalent in regions that have historically experienced more intense drought. These SNPs are putatively adaptive or physically close to putatively adaptive sites within the genome. (D) GEA of Baseline samples (X) reveals 10 Kbp windows associated with historic winter precipitation (1980-2009). Dashed line indicates the Bonferroni-corrected significance threshold. Pink dots represent peak 10-Kbp windows that were selected for further analyses (see Supplemental Methods for SNP set selection and Fig. S3-4 for all other climate variables). (E) Frequency changes over time, through extreme drought, for 215 climate-associated SNPs within Time Series samples (triangles) from a southern population (see Fig. S6 for all other populations). Each line represents a single regression (one per locus) whose slope is the response to selection (S). (F) Distribution of the strength of selection (S = slope of SNP change) across the climate-associated SNPs in Time Series samples (triangle) for this southern population (see Fig. S7 for all other populations). Histograms summarize the population of slopes depicted in panel E and give the number of observed SNPs within 0.1 S bins. Positive S indicates positive directional selection toward the heat/drought-associated allele identified during baseline GEA (i.e., pre-drought spatial associations of climate and SNPs). The red vertical line gives the median S for this population. (G) Distribution of response to selection observed in 1000 permutations of non-climate associated SNP in Time Series samples (triangles) for the same southern population, shown as grey histograms (see Fig. S9 for all other populations). Median response to selection from climate-associated SNPs (red vertical line, as in (F); note the difference in scale of x-axes) is compared to the median of the permuted distribution (grey vertical line). In this population, the observed response to selection at climate-associated SNPs exceeds that of the random permutations in this population, suggesting that allele frequency change is driven by selection instead of drift.

Next, to assess evidence for rapid evolution in response to exceptional drought, we tracked the changes in frequencies of climate-associated SNPs through the drought by sequencing a time series dataset composed of 11 populations sampled annually between 2010 and 2016 (N= 401 individuals; Fig. 2B, Table S5). We hypothesized that <u>the frequency of alleles associated with drought in the GEA should rise through time in each population if the contemporary extreme drought imposed similar selection as</u> exerted in <u>historically</u> drier and hotter places. We tracked frequency change for the 215 unlinked SNPs that monotonically change with climate (Table S4; Supplemental Methods), with alleles associated with drier or warmer historical conditions coded as 1. For each SNP, we defined response to selection (S) as the slope of a generalized linear binomial model for SNP frequency over time (Fig. 2E; Fig. S6 (*11*)). Since frequency = 1 is the drought-associated state of each SNP, a positive S is consistent with selection for increased adaptation under drier or warmer conditions. This approach reveals extensive evidence of changes in frequencies of climate-associated SNPs across the range of *M. cardinalis* (Fig. 2E,F; Fig. S6-7). Median selection coefficients were significantly different from zero (the benchmark for nochange) in 5 out of 11 populations (Table S6). Many individual climate-associated SNPs experienced weak positive selection (S=0 to 0.2), with a considerable number of SNPs showing moderate (S=0.2 to 0.5) or strong selection (S>0.5) towards the drought-associated state. Some alleles also experienced negative selection, particularly in northern populations (Fig. S6-7), with four of seven central and northern populations showing median selection away from the direction predicted by SNP-environment associations (i.e., decreases in the drought-associated allele). This suggests that not all SNPs associated with historic adaptation to climate will be important in future adaptation to climate. In common gardens comparing morphological, biochemical, and phenological traits of resurrected ancestors (families sampled pre-drought) and descendants (families sampled throughout the drought), southern and northern populations also differed in the magnitude and direction of trait evolution (*18*). Populations with positive median S tended to evolve greater rates of photosynthesis (b = 0.37±0.15, p = 0.04) and lower rates of stomatal conductance (b = −0.34±0.08, p = 0.005), while populations with negative median S evolved lower ates of photosynthesis and higher rates of stomatal conductance (Fig. S8, Table S7). Thus, populations with positive selection for heat/drought alleles evolved greater carbon capture relative to water loss, while populations with selection against heat/drought alleles evolved lower carbon capture relative to water loss. These seemingly maladaptive trajectories of the latter populations could be due to mismatched germination cues or trade-offs with unmeasured traits.

In addition to selection imposed directly or indirectly by the drought, genetic drift may also have driven allele frequency change, particularly because population sizes were small and usually declining during drought. To estimate the extent to which allele frequencies could be expected to change under drift alone, we created a null distribution of change in allele frequency by randomly selecting non-climate-associated SNPs and calculating their slopes of change over time (Supplemental Methods). As expected under neutrality, null distributions of mean S for non-climate-associated SNPs were centred on zero (with standard deviations ranging from +/-0.21 to +/-0.51 across populations). While all populations had positive selection for the drought-associated state in some SNPs, three populations had median selection strengths significantly greater than a null distribution of median selection strength from non-climate associated SNPs (Table S6, Fig. 2G, Fig. S9; Populations 3, 4, 11). Non-random selection for alleles associated with hot and dry environments during 2012-15, a time period of consistently high drought severity (Fig. 1B, Fig. S2), suggests rapid evolution by natural selection during extreme drought.

So far we have demonstrated two parts of evolutionary rescue. *Mimulus cardinalis* populations were driven into decline during drought, and some populations rapidly evolved non-random increases in historically drought-associated alleles, though to different extents. To fully demonstrate evolutionary rescue, we must assess if populations recovered and if rapid evolution was responsible for this recovery. To test for recovery, we again used field demographic data on survival, growth, reproduction, and recruitment to infer *r* via IPMs, this time using post-drought observations from the 2015-16 to the 2017-18 annual intervals (Fig. 3A, Supplemental Methods), which is the relevant time window given life-history lags (*24*). Many *M. cardinalis* populations showed evidence of post-drought recovery (Wilcoxon rank-sum test: V=117, p=0.009; Fig. 3B, S1B, D; Table S1), with mean growth rates that were above replacement (*r*>0) in 14 out of 19 populations. Still, there was much variability in the magnitude and stability of the recovery, with some populations showing a downward trajectory and three populations remaining locally extinct (populations 14, 15, 17) within our monitoring area (Fig. 3B, Fig. S1B, D).

**Fig 3.**
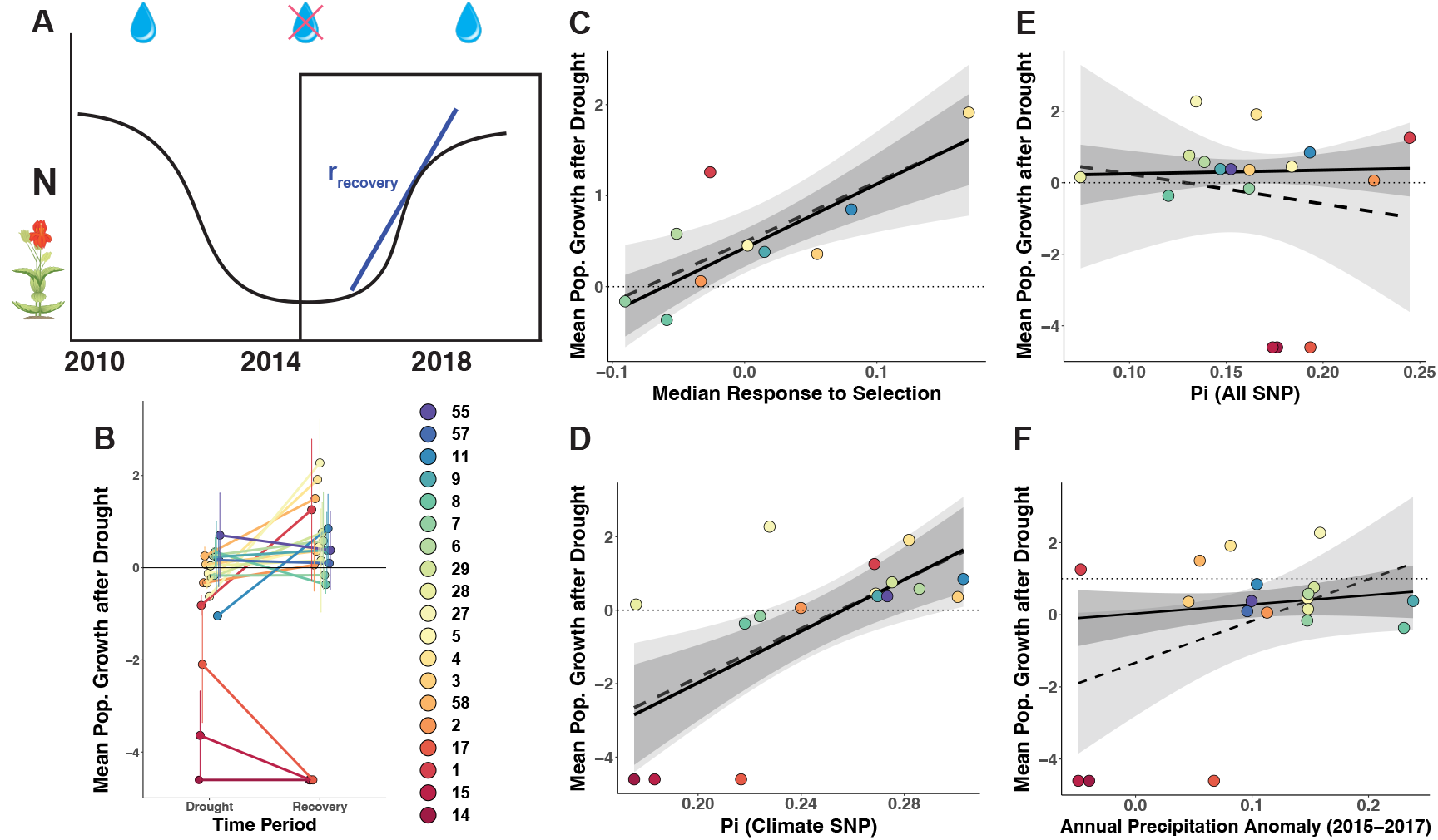
Rapid evolution and nucleotide diversity explain demographic recovery. (A) Recovery phase of evolutionary rescue, indicated by the rectangle. r_recovery_ represents mean population growth rate (*r*) during the drought time period (2015-16, 2016-17, 2017-18). (B) Mean *r* during drought (2012-13, 2013-14, 2014-15) and recovery time periods shown as latitudinally colored points with lines connecting consecutive values for a given population. The horizontal black line depicts rate of replacement (*r* =0); values below this line indicate a population that is projected to decline. (C) Greater median response to selection during severe drought predicts mean population growth rate during recovery. Ten populations have both demographic estimates and time series genomics. (D) Having greater nucleotide diversity (π) for climate-associated SNPs prior to drought onset predicts greater mean population growth rate during the recovery period. Sixteen populations have both baseline sequencing and demographic estimates. (E) Genome-wide nucleotide diversity (π) across all SNPs does not predict mean population growth rate during recovery. (F) Population recovery after severe drought (2015 to 2018) is not explained by less anomalous winter precipitation (or other climate variables; Table S2). For C to F, each point represents one population, with rainbow colors depicting latitude of origin (blue = northern populations, red =southern populations). Population codes are given for each population (Table S3). Solid lines and darker shading depict predicted slopes and their 95% confidence intervals from robust regression. Dashed lines and lighter shading depict predicted slopes and their 95% confidence intervals from ordinary least squares regression. Dotted horizontal lines depict rate of replacement (*r* =0), above which populations are projected to increase and below which populations are projected to decline.

Variation in post-drought population demography was strongly explained by response to selection at climate-associated loci during drought, providing evidence consistent with evolutionary rescue (Fig. 3). Among the subset of populations for which we had 2010-2016 timeseries genome sequences and suitable demographic recovery data (Table S3), greater rapid evolution (measured as greater median positive selection on the climate-associated SNPs) predicted greater mean population growth rates (Fig. 3C; ordinary least squares regression (OLS): P=0.011, R^2^=0.52;robust regression: P=0.012; Table S7). Furthermore, nucleotide diversity of climate-associated SNPs prior to drought onset predicted greater population recovery (Fig. 3D; OLS: P=0.005, R^2^=0.38; robust regression: P=0.004; Table S7). Critically, SNPs identified by GEA were necessary for explaining demographic recovery; genome-wide nucleotide diversity was uncorrelated with post-drought population growth rate (Fig. 3E; OLS: P=0.55, R^2=-^0.04; robust regression: P=0.86; Table S7).

An alternative explanation for population recovery could be ecological rescue, where populations recovered simply because the drought ended. To assess ecological predictors of population recovery, we calculated post-drought climate anomalies as the deviations from 30-year pre-drought climate averages (1980-2009) for the means of each of eight precipitation and temperature-related variables during the recovery period (2015-17; Supplemental Methods). If demographic recovery is attributable simply to cessation of drought, then populations with wetter and cooler post-drought climate anomalies should have higher population growth rates. However, climate anomalies rarely and only weakly predicted mean population growth rate following the drought (Fig. 3F, Table S2). Another alternative explanation for population recovery could be negative density-dependence, where the populations with the greatest decline had greatest scope for recovery following the drought; however, we did not find a negative correlation between mean population growth during vs. after drought (Fig. S10; r=0.14, P=0.57), which suggests population recovery is not due to density-dependent demographic compensation. Further, it is improbable that density-dependence could account for the relationships between post-drought population recovery and genetic metrics (i.e., selection on, and standing nucleotide diversity of, climate-associated SNPs). Thus, currently our best explanation for the observed variability in population recovery is differences in capacity for evolutionary rescue.

Combining demographic and genomic data across space and time, we observed all three requirements for evolutionary rescue in *M. cardinalis (4, 5). Mimulus cardinalis* populations declined during severe drought. Rapid evolution of drought-associated alleles occurred during this population decline. Rapid evolution then predicted differences in demographic recovery. Furthermore, populations harboring higher levels of nucleotide diversity at climate-associated SNPs prior to the drought also had higher population recovery after drought, which implicates adaptation from standing variation. Together these results show the importance of standing genetic variation and evolution to drought resilience. Given the severe and far-reaching impacts of the Western North American megadrought (*25-27*), significant pressure to rapidly evolve has likely been widespread (*18, 28-30*). We have shown that evolutionary rescue is possible in a relatively short-lived species (although longer-lived than most focal organisms in resurrection studies and studies of evolutionary rescue) with high genetic variation, despite small census population sizes. These results remain to be validated in species of conservation concern that may have much lower genetic diversity and longer generation times. It is possible that evolutionary rescue, which we have demonstrated here for *M. cardinalis*, could have also occurred in other short-lived species, and might be critical for long-term population viability. Indeed, prior work on evolutionary rescue in a microbial laboratory experiment suggests that populations and communities exposed to strong but non-lethal stress may be better able to survive even more severe events in the future (*30, 31*). Thus, populations of *M. cardinalis* and potentially other organisms may be better placed to survive future climate extremes given the significant drought exposure. Alternatively, southern populations might be nearing physiological limits (*32*) and selection might deplete genetic variation and lessen future response to multivariate climate stressors.

The ability of genomics to predict future demographic outcomes is one of the most important open questions within conservation genomics. There is considerable debate on the importance of genome-wide diversity in conservation, where some studies find genome-wide diversity predictive of population resilience (*33, 34*) while others find that direct information on adaptive genetic variation is needed (*35, 36*). Moreover, validating the function and adaptive utility of GEA-identified loci remains challenging (*37, 38*). Greater investigation into the ability of climate-associated nucleotide diversity to predict demographic outcomes of systems undergoing climate change may help assess broader utility of sequencing efforts for conservation management. Within such efforts, we have shown that genotype-environment association is critical to identifying the subset of genetic variation best able to predict demographic resilience to climate change when clear climatic stressors are known (Fig. 3D vs. 3E). This provides a unique validation of GEA inferences with independent demographic data, while lending support to the need to identify adaptive loci within the genome. However, these GEA-identified SNP are not necessarily indicative of the many other environmental stressors that populations will face. Thus, we recommend using GEA-identified SNP sets, alongside genome-wide diversity approaches, when using nucleotide diversity to inform conservation management efforts for population resilience.

These efforts could be used in conjunction with other spatially-based genetic diversity metrics such as the mutation-area relationship (*39*) and metrics of genomic vulnerability to climate change (*35, 36, 40, 41*). Ultimately, space-for-time approaches, based on the idea that genetic variation spread across space can be used to predict genetic variation needed across time, are broadly used in evolutionary biology and conservation genetics (*37*). Our study supports the utility of space-for-time approaches in predicting real world eco-evolutionary outcomes.

## Acknowledgements

J. Paul, A. Campbell-Craven, P. Beattie, J. Smith, A. Wilkinson, B. Econopouly, C. Fallon, E. Hinman, M. Bayly, D. Picklum, J. Perce, A. Agneray, M. Bontrager, M. Bayly, Q. Li, B. Gass, L. Super, E. Okun, T. Usui, R. Wilson, M. Nagaraj, and J. Wynne aided in demographic monitoring and seed collections. H. Branch, J. Zajonc, K. Saller, K. Beckett, and C. Longan aided in seed germination and greenhouse management. C. Elphinstone, and W. Cheng aided in DNA extractions and ordering. J. Whitton provided lab space and equipment. L. Fishman provided the *M. cardinalis* genetic map. S. Heredia provided illustration and figure design. Genome Quebec carried out library prep and Illumina Sequencing. Advance Research Computing at UBC provided computational resources and support. Special thanks to D. Lowry, W. Wetzel for advice and mentorship. Angert, Lowry and Wetzel labs provided feedback. **Funding:** This work was funded by a Genome BC Grant to A. Angert & L. Rieseberg; NSERC Banting, NSERC PDF, Killam Fellowship, and PRI Fellowship to D. Anstett; and NSERC Discovery grants to A. Angert and L. Rieseberg. S. Sheth was supported by NSF DEB-2131815 and USDA National Institute of Food and Agriculture Hatch 7002993. S. Sheth and A. Angert were supported by NSF DEB-2311414. **Authors contributions:** D.N.A. led the project and was involved in all aspects. D.N.A., A.L.A, and L.H.R. conceived the project. R.J., S.N.S. and J.A. provided conceptual support. J.A. and D.N.A. performed bioinformatic and statistical analyses with support from M.J., K.H., J.M.L.G., R.J., A.L.A., and L.H.R.. A.L.A., S.N.S. and D.N.A. conducted demographic field work and seed collection. A.L.A. and S.N.S conducted demographic analyses. D.N.A., and A.L.A. curated seeds and generated leaf tissue for DNA extraction. D.R.M created the DNA extraction protocol. D.R.M & D.N.A conducted DNA extractions with support from M.T.. D.N.A., A.L.A., J.A., S.N.S, D.R.M. wrote the paper. H.A.B. contributed phenotypic analysis and guidance on their biological significance. All authors provided edits and feedback. **Competing interests:** Authors have no competing interests. **Data and materials availability:** All code is available on GitHub (https://github.com/anstettd/evol_rescue) and archived on Zenodo (*42*). SNP tables and additional supplemental files are also available on Zenodo (*43*). Raw reads have been deposited in the NCBI Sequence Read Archive under BioProject PRJNA1018529.

