## Supplemental Material for "Evolutionary rescue during extreme drought"

**Supplemental Material for Anstett et al.**

### Methods

#### *Study System*

*Mimulus (Erythranthe) cardinalis* (Douglas ex Benth.) is a short-lived perennial forb found along streambanks and seeps from Southern Oregon, USA, throughout California (USA) and into Northern Baja California, Mexico (*44*). *Mimulus cardinalis* is hummingbird-pollinated and outcrossing but also self-compatible. It propagates through seed and vegetatively through rhizomes. The plant grows along an aridity gradient with reliably higher precipitation in Oregon, to lower and relatively more variable precipitation and higher temperatures in Southern California. This aridity gradient is associated with variation in physiology and phenology, with faster life history (*3*) and greater growth and photosynthetic rate (*17*) in southern populations. The growing season of *M. cardinalis* is characterized by declining soil moisture following the spring freshet, making this plant a good indicator of the effects of drought.

#### *Population Demography*

Since 2010, we have conducted annual demographic censuses and plant measurements in 21 wild populations spanning the species' latitudinal range. Here we used these data to estimate *per capita* population growth rates from just prior to drought onset through the peak of the drought. Full methods are available in Sheth and Angert (*2*). In brief, populations were visited during August-October, after most reproduction was complete for the growing season. From these records, we derived *per capita* annual survival (0 or 1), size (total stem length), growth (annual change in stem length), probability of flowering (0 or 1), fertility (number of fruits per plant times population mean seed number per fruit), and recruitment (the proportion of seeds from the previous year that became new plants). In total, the fates of 20,353 plants were recorded from 2010 through 2019 (range: 179–2,101 individuals per population; mean = 1,018 individuals) across sampling plots that average 471 $m^2$ (range 41 – 4800 $m^2$ due to spatiotemporal differences in habitat structure and plant density).

To estimate the projected asymptotic population growth rate ($\lambda$) for each population, we used Integral Projection Models (IPMs) (*19*). We constructed global models of each of four vital rates (survival probability, growth, flowering probability, and fruit number) as functions of size (ln-transformed total stem length in year *t*, fixed effect), year (random effect), and population (random effect) and then extracted population- and year-specific coefficients. Survival and flowering probabilities were modeled with binomial distributions, growth was modeled with a Gaussian distribution, and fruit number was modeled as a negative binomial distribution without zero inflation (the latter of which was chosen after comparing to a Poisson distribution with and without zero inflation). We explored various random effects structures where years and/or populations could have different slopes and/or intercepts with respect to size and selected the model with the lowest Akaike Information Criterion that did not give singularity warnings. Final vital rate models, including random effects structures, are provided in Table S9, and population- and year-specific coefficients used to parameterize each vital rate model are in Table S10. We used the fitted vital rate models to build a discretized matrix for each population-year with 100 size bins, ranging from 0.9 times the minimum to 1.1 times the maximum size observed in each population, correcting for the "eviction" of individuals falling beyond this size range by assigning individuals to the smallest size bin in the case of offspring and to the largest size bin in the case of large adults (*45*). We then calculated $\lambda$ as the dominant eigenvalue of the population- and year-specific discretized matrix (Table S10). Because of right-tailed skewness, we log-transformed $\lambda$ (after adding a small value of 0.5 to account for zeros). These log-transformed values are also known as the intrinsic rate of population growth, *r*. We also estimated generation time for each matrix as the expected age at reproduction for a cohort using the 'cohort' method of the 'gen_time' function of the Rage package (*46*), resulting in a mean generation time estimate of 3.92±0.22 years (Table S1).

We described demographic trajectories during the drought as the mean *r* during the core drought period (2012-13, 2013-14, 2014-15) (Table S1). This metric captures differences among populations in the absolute severity of the drought on projected population persistence, where a mean below 0 indicates a population that is projected to decline to extinction. Note that *r* is a projection of the *per capita* rate of change in *N* over time, making this metric a good proxy for the slope of population decline depicted in Fig. 1A & slope of population recovery depicted in Fig. 3A. Analogously, we described demographic trajectories after the drought (Fig. 3A) as the mean *r* during the post-drought recovery period (2015-16, 2016-17, and 2017-18) (Table S1). This sampling period allowed enough time to reflect time lags in response to selection due to perennial life history (*24*). We derive metrics of demographic trajectories based on *r*, rather than simple population size counts, because *r* captures differences in population dynamics when individuals of different sizes have different vital rates. For example, a population of 100 post-reproductive, senescent individuals has a different growth potential than a population of 100 juveniles with their full reproductive life span ahead of them. This is the fundamental reason for using structured population models (where structure is defined by the state variables that predict demographic vital rates. Common state variables are age or, in our case and the case for most plants, size). Thus, intrinsic growth rate from a structured projection, rather than a simple count-based metric of density or population size, better captures a population's demographic capacity to expand or maintain its size in future years.

One population (Mill Creek) was excluded because all plots washed out between the 2010 and 2011 censuses, and flash floods precluded population access in 2013, causing transition estimates from 2010-11, 2012-13 and 2013-14 to be inestimable. Plot relocation to a more stable area of the drainage in subsequent years confounded space and time, hence this population is removed from all analyses. Another population (Deer Creek) was excluded from demography estimates because annual plot wash-outs caused individual relocation and identification to be unreliable; this population was included in genomic analyses but removed from all analyses involving *per capita* demographic estimates. We performed all analyses in R 4.2.1 using code modified from Sheth and Angert (*3, 47*).

*Seed & Tissue Collection*

A "Baseline" seed collection was established prior to the onset of the 2012-2015 extreme drought. Seeds were collected from 55 populations across the range of *M. cardinalis* from 2007 to 2011 (Table S2). Individuals were sampled at least 1m apart to capture unique genetic individuals. Seeds were germinated in a greenhouse, where young expanding leaves were collected and frozen for tissue extraction. We collected seed pods from up to 10 individuals per population. A "Time Series" seed collection was also carried out for 12 populations throughout the range of *M. cardinalis* (Fig. 1; Table S2). Seed pods were collected yearly between 2010 and 2016. Up to 10 plants from each population were sampled, with some temporal gaps due to fire, population inaccessibility or lack of viable seed. For Time Series plants, we carried out a refresher generation crossing within population/year as outlined in Anstett, Branch and Angert (*5*). Leaf tissue was collected for sequencing for each refreshed line. In total, 655 individuals across Baseline and Time Series collections were selected for DNA extraction and sequencing. 402 individuals were present in the Time Series collection and 347 individuals were in the Baseline collection, with 94 samples shared between the Baseline and the Time Series data sets.

*Climate Data*

Climate data were downloaded from Climate NA (*22*). We extracted nine climate variables associated with drought: MAT = Mean annual temperature (°C); MAP = Mean annual precipitation (mm); PAS = Precipitation as snow (mm) between August in previous year and July in current year; EXT = Extreme temperature over 30 years; CMD = Hargreaves climatic moisture deficit (mm); Tave_wt = Winter mean temperature (°C); Tave_sm = Summer mean temperature (°C); PPT_wt = Winter precipitation (mm); and PPT_sm = Summer precipitation (mm). We extracted 1980-2009 averages for each population. We also extracted yearly data (PAS), seasonal data (Tave_wt, Tave_sm, PPT_wt, PPT_sm) and monthly data (MAT, MAP, CMD) from 1979 to 2019. Climate during the 2012-2015 drought period and 2015-2018 recovery period were calculated as an average for each of the listed years. To calculate climate anomalies, we subtracted 30-year average data for the period 1980-2009 from each of 2010 to 2019 data for each climate variable. Then anomaly averages were calculated across years for 2012-2015 for population decline and 2015-2018 for population recovery. Precipitation values were logged to express them as proportional changes instead of absolute changes, since sites differ dramatically in overall precipitation. Water year (October previous year to September current) was used for MAT, MAP and CMD, while seasonal and PAS data already included an appropriate monthly range to reflect the life cycle of *M. cardinalis*. These climate anomalies were then regressed against mean lambda for population decline (2012-2015) and recovery (2015-2018) using the lm command in R.

*DNA Extraction*

DNA was extracted following a modified method of Xin & Chen (2012). 100 ug/mL Proteinase K was added to the CTAB Lysis buffer to reduce enzymatic oxidation; centrifugation speeds were increased to 6000 rcf following DNA-CTAB complex precipitation. The MagAttract beads were suspended in 55 µL of TE Buffer (10 mM TRIS, 0.1 mM EDTA, pH 8.0) and incubated at 37 ºC, 150 rpm for 30 minutes. 54 µL of the sample was transferred to a clean PCR plate. Purity was accessed using 2 µL of DNA elution on a Nanodrop 2000 (Thermo Scientific™), and DNA concentration was quantified with the Qubit 2.0 Broad Spectrum kit (Invitogen™) using 2 µL of DNA elution. The full-length protocol can be found at protocols.io/private/0E020E942EA511EB83910A58A9FEAC2A?step=22.

*Library Prep and Sequencing*

A minimum of 150 ng DNA per sample was submitted for WGS library preparation and sequencing. Both NEB Ultra II Shotgun DNA library preparation and sequencing were conducted by the Centre D'expertise et de Services Génome Québec. The libraries were sequenced at 150-bp PE using the Illumina NovaSeq 6000 system.

*Preprocessing Raw Reads*

Variants were called on the 655 individual gnomes that were sent for library preparation sequencing. Sequences were trimmed for low quality reads using trimmomatic (v0.39) with standard settings and ILLUMINACLIP adaptors_illumina.fasta:2:30:10:8:T SLIDINGWINDOW:4:15 MINLEN:36 (*48*) (*7*), and aligned to the *M. cardinalis* reference genome, CE10_v2 (www.mimubase.org) using bwa-mem (v0.7.17-r1188) (*49*), and indexed with samtools(v1.6) (*9*) with standard settings. PCR duplicates were marked with picard MarkDuplicates (v2.25.0) (*50*). Read groups were then added to the marked bam files with gatk AddOrReplaceReadGroups (v4.9.1).

*Variant Calling for Gold Dataset*

To perform variant calling, we followed the best practices recommendations of the Genome Analysis ToolKit (GATK) *(51)* and executed steps documented in GATK's germline short variant discovery pipeline (for GATK v4.2.0.0). We conducted the workflow on the twelve samples that had the highest coverage (16-21X) in order to recalibrate the base quality scores for the full dataset, and designated this as the "gold dataset". First, HaplotypeCaller was individually run on the marked and sorted bam files for each of the gold dataset samples to produce genomic VCFs (g.vcf or GVCFs). Then, the twelve GVCFs were consolidated then split into eight GenomicsDB datastores, one for each chromosome using GenomicsDBImport (v4.2.0.0). We then used GenotypeGVCFs to conduct joint genotyping and created a GCVF for each chromosome. Next, we used SelectVariants to generate separate VCF files for SNPs and indels. The SNPs and indels for each chromosome were then hard filtered with VariantFiltration using the following parameters: mapping quality (MQ) >50.0, AN =21.6 (12*2*0.9), −1.0> strand odds ratio (SOR) <1.0, >0.25 Minor Allele Frequency <0.75, excess heterozygosity >10.0, −1.0> BaseQRankSum < 1.0, and depth of coverage within one standard deviation from the mean for each chromosome for both SNPs and indels. The final variants were selected with SelectVariants, and the genotype information was stripped with MakeSitesOnlyVcf. The final chromosomal VCF files were then merged into two single VCF files, one for SNPs and one for indels using GatherVcfs, and indexed with IndexFeatureFile.

*Variant Calling with Base Recalibration for Full Dataset*

The final SNP and indel VCF files from the gold dataset were used to recalibrate the base quality scores of the sorted and marked BAM files for the full dataset using BaseRecalibrator and ApplyBQSR. Haplotype calling was then conducted on the recalibrated BAM files with

HaplotypeCaller. The resulting GVCFs were consolidated then split into GenomicsDB datastores. Since 655 samples were consolidated, the datastores were split into 5,000,000 genomic intervals for each chromosome, producing 86 datastores. We then used GenotypeGVCFs to conduct joint genotyping and created a GCVF for each datastore. The GCVFs were then merged into eight GVCFs, one for each chromosome, and indexed using GatherVcfs and IndexFeatureFile. Next, we used SelectVariants to generate separate VCF files for SNPs and indels. We then built a recalibration model to score variant quality for filtering purposes with VariantRecalibrator for both SNPs and indels using the gold dataset as the truth dataset and the following parameters: prior=10.0, -an MQRankSum -an ReadPosRankSum -an FS -an MQ -an SOR -an DP –trust-all-polymorphic --truth-sensitivity-tranche 100.0 –truth-sensitivity-tranche 99.0 –truth-sensitivity-tranche 90.0 –truth-sensitivity-tranche 70.0 –truth-sensitivity-tranche 50.0, and –max-gaussians 6. The SNPs and indels for each chromosome for all individuals were then filtered with ApplyVQSR, using a truth sensitivity filter level of 50.0 using the raw and recalibrated/filtered VCF files. We then gathered the SNP VCFs into a single VCF using GatherVcfs and indexed with IndexFeatureFile. The final set of SNPs were selected using SelectVariants with the following parameters: restrict alleles to biallelic, > 0.01 allele frequency < 0.99. Baseline and Time Series data were separated into different vcf files using SelectVariants.

*Genotype-Environment Association*

Genotype-environment association (GEA) was carried out using both a SNP-based and a window-based approach. The SNP-based approach accounts for population structure, while the window-based approach accounts for genome structure. The association of all SNP allele frequencies within the baseline dataset to the 9 climate variables were assessed using BayPass ver. 2.2 (*20*). Population structure was taken into account randomly sampling 10,000 putatively neutral SNPs and using them in the BayPass core model (*20*). Bayes factors were calculated under the standard covariate model to evaluate association with 30-year average climate data from the 9 aforementioned climate variables. SNPs with a high threshold of -log10BF > 30 were considered climate-associated SNPs. The -log10BFs from all SNPs were then employed for the window-based approach WZA that uses supporting information from adjacent SNPs to establish regions most strongly associated with each climate variable (*21*). Using 10,000 non-overlapping bp windows, we calculated WZA scores for 38,632 windows. Empirical p-values were generated for each window. We set a Bonferroni significance threshold of $0.05/38,632 = 1.294264 \times 10^{-6}$. For all windows above this threshold, we selected the peak (top) window within each series of adjacent windows that were above the threshold. When two windows were extremely close to each other (<0.3 log P-value units away from each other) we kept both windows rather than making a somewhat arbitrary choice. We then carried out a double filtering approach where SNPs with -logBF >10 from significant windows were considered to be climate-associated. These SNPs were combined with the -logBF > 30 set from BayPass, resulting in a total of 605 climate-associated SNPs used in downstream analyses.

*Nucleotide Diversity*

Missing SNP genotypes were imputed using Beagle 5.1 with default settings. A genetic map was used from (*52*) with physical/genetic relationships from (*53*). To further account for linkage across these SNPs, we binned the 605 climate-associated SNPs using the *clump* command in PLINK (*23*).

Each SNPs was clumped within 250 kbp and r2 > 0.4 into a single LD block. The SNP with the highest Bayes Factor (from BayPass) was then selected for each clump, leaving 341 unlinked SNPs. Per-SNP nucleotide diversity ($\pi$) was calculated using vcftools. Average $\pi$ was calculated for all SNPs across the entire genome (without invariant sites) and for the set of 605 climate-associated SNP for all 55 Baseline populations. Both of these metrics differ from traditional estimates of genome-wide PI, which is typically calculated using all invariant sites as well. Hence our metric will have a greater magnitude than those typically reported.

*Defining Climate-Change-Associated Alleles*

Allele frequencies were calculated for each of the 55 Baseline populations and each Time Series population/year combination (62 unique "populations"). To facilitate interpretation, we defined allele frequencies of all climate-associated SNPs to ensure 1 always reflected fixation of the drought-associated (decreased precipitation, increased temperature, increased moisture deficit) allele and 0 reflected fixation of the non-drought-associated allele. This was done by associating the frequency of each SNP from each of the 55 Baseline populations with 30-year historical climate using glm(), family=binomial and filtering the SNPs to include only those loci with monotonic associations between SNP frequency and baseline climate; to this end, we filtered out SNPs with a slope of frequency vs. climate <0.3 on the logit scale. We also removed all SNPs with an average frequency across baseline populations of <0.03 or >0.97 because these SNPs also lacked a strong monotonic relationship with baseline climate (regardless of SNP frequency change across time). In total this filtering removed 231 SNPs, leaving 374 SNPs. To further account for linkage across these SNPs, we again binned using the *clump* command in PLINK (*23*), yielding 215 SNPs for temporal analysis.

*Temporal Selection Analysis*

We described the change in frequency of all 215 unlinked, monotonically climate-associated, biallelic SNPs over time from 2010 to 2016 using logistic regressions (glm function with family=binomial in lme4 package), across each of the 11 Time Series populations, carrying out one regression per SNP, per population. Population/year combinations did not have equal sample sizes due to variability in the number of available seed lines (*5*). Thus, to ensure equal weighting of individuals, we transformed the abundances into individual 0's and 1's for the purposes of the regression. The logit-transformed slope of each of these binomial regressions gives an estimation of the selection coefficient (S) during the extreme drought *(19)*, with S>0 suggesting positive selection for the drought-associated allele and S<0 suggesting negative selection against the drought-associated allele. A small number of SNPs were identified by more than one environmental variable. In those cases the S was identical, since the year and frequency data were the same, and we only included one S value for each SNP. Extremely high selection coefficients were produced for SNPs with poor sampling (e.g., multiple years not represented at a particular SNP/population). These SNPs also had a very high standard error (SE); thus, we removed all regressions with SE>5.

To further visualize the S distribution, we plotted histograms of selection coefficients per population for the monotonically climate-associated SNP set using 0.1 S bandwidths. We also calculate median S across the selected climate-associated SNPs for each population. We carried out a Wilcoxon signed rank test to determine if the medians for each population were significantly greater than 0.

To identify if median S is greater than would be expected by genetic drift alone, we estimated a null distribution of SNPs that were not climate-associated. This was done by randomly selecting an equal number of SNPs that are not correlated to any of our 9 focal climate variables as established by BayPass (-log10BF<0). For consistency with the handling of climate-associated SNPs, we again removed all regressions with SE>5. This process was carried out 1000 times, generating 1000 distributions of S for non-climate-associated SNPs. To avoid artificially biasing this distribution in favor of the major or minor allele, we randomly assigned the major or minor allele to each non-climate-associated SNP prior to randomly selecting SNPs. Major and minor alleles were defined using the Baseline genomic data set.

To test for a difference between climate-associated and non-climate associated medians, we calculated a permutation p-value using the distribution of medians from the 1000 non-climate-associated SNP distributions. This tests the hypothesis that S for climate-associated SNPs is significantly greater than the null distribution, which is likely driven by drift alone.

*Phenotypic trait evolution*

We reanalyzed data from Anstett et al. 2021, which quantified rates and magnitudes of trait evolution during the drought in a resurrection common garden with the same populations and year ranges as the current study. The prior study measured leaf-level gas exchange (photosynthetic carbon assimilation and stomatal conductance), leaf morphology (specific leaf area and leaf water content), and phenology (day of first flower) on seed families collected before, during, and after the drought. Here we quantified rates and directions of trait evolution for each population as its slope of trait value versus family year of origin within the dry watering treatment (the relevant selective environment). These population-specific slopes were extracted from linear mixed models for each trait modeled as a function of year, population, watering treatment, and their interactions, with random intercepts for seed family and randomized block on the greenhouse benches. All variables were standardized to a mean of zero and a standard deviation of one prior to analysis. We then used multiple regression to relate slopes of trait evolution to our index of genomic selection, median S. We constrained the maximum number of predictors to two, based on population number, resulting in 3 subsets of predictors organized by biological function: gas exchange (photosynthesis and conductance), morphology (specific leaf area and leaf water content), and phenology (flowering time). Model selection across all possible one- and two-variable models (i.e., not grouping predictors by biological function) identified the two-variable model with both gas exchange variables as the top model (lowest Akaike Information Criterion score; results not shown). Predictions from the gas exchange model were visualized using the ggeffects package (*54*). Predictions for each focal predictor were estimated at the mean of the non-focal predictor using the margin = “mean_reference” command.

*Testing for evolutionary rescue*

We tested whether mean population growth rate (*r*) during recovery (2015-16 to 2017-18) was associated with median response to selection on climate-associated SNPs using a simple linear model and assessed significance with type III anova with the *lm()* and *Anova()* commands in R (*55, 56*). We also used simple linear models to test whether recovery population growth rate was predicted by pre-drought nucleotide diversity ($\pi$), both genome-wide and when restricted to the subset of all 341 selected climate-associated SNPs. For response to selection, 10 out of 11 populations were included (Table S2), since reliable demographic data for Deer Creek were not available as explained previously. For $\pi$, we included 17 populations out of the 19 demography populations (Table S2) because the populations Canton Creek and South Fork Middle Fork Tule River did not have Baseline sequencing data. We visualized these relationships as regressions in scatter plots using *ggplot2 (57)*. To rule out ecological rescue, we associated climate anomaly with population growth rate for all 17 demography populations (as outlined in the Climate Data selection) and we calculated the Spearman rank correlation between population growth rates during and after drought. We failed to detect a significant effect for any climate variable (Table S1), and hence did not include the effect of climate as a covariate in relationships between demographic recovery and genomic metrics. To assess the effect of outliers, we compared relationships estimated from ordinary least squares regression (lm function of base R) and robust regression (rlm function of MASS package (*58*)) (Table S8). Significance of slopes estimated from robust regression was obtained with a robust F test (f.robftest function of the sfsmisc package (*59*)).

*Subset of Climate Variables*

To test the impact of climate variable selection, we recalculated methods downstream from GEA using a subset of environmental variables. Rather than the 9 variables in the full analysis we selected only SNPs associated with the three environmental variables, CMD, Tave_sm, and PPT_wt, that best predicted decline in growth rates during the drought (Table S1), while using the same SNP set filtering methods outlined in above sections. This yielded 103 SNPs for the temporal selection analysis, and 159 SNPs for the Pi analyses. As before, we describe the change in frequency of the selected SNPs over time from 2010 to 2016 using the slope of logistic regressions, defined as Response to Selection (S), calibrated so positive values indicate evolution towards a climate change adapted state. A distribution of randomly selected non-climate associated SNPs were again generated per population and used as a significance test for each median S value (see Temporal Selection Analysis section). S and Pi were then used to explain mean population growth rate after drought.

### Supplemental Results

The results from the analysis using the subset of climate variables were qualitatively similar to the analysis with all 9 climatic variables. For each population we again observed a distribution of values of S ranging from just under -1 to just over 1, with Median S being similar for most populations (Fig. S11). Critically, population 3, 4 and 11, and additionally 1 as well, show similar positive values for median S, while 7,8 and 10 show negative values. Under this analysis, fewer populations were statistically different from drift (Table S11, Fig. S12), likely due to decreased power of considering fewer SNPs. Populations with greater median response to selection (from the SNP set generated with only CMD, Tave_sm, and PPT_wt associated SNPs) had higher mean population growth rate after drought (p=0.0006, $R^2$=0.76, robust regression =0.001; Fig. S13A). Populations with greater Pi (again from the reduced climate SNP set) also had greater mean population growth rate after drought (p=0.004, $R^2$=0.68, robust regression =0.009; Fig. S13B). Overall, similar conclusions can be drawn when using the subset of climatic variables that were associated with demographic decline versus using all nine climatic variables. For the climatic subset there is greater variation explained and greater statistical support for rapid evolution explaining population recovery, but lower support for S being different from drift.

### Supplemental Figures & Tables

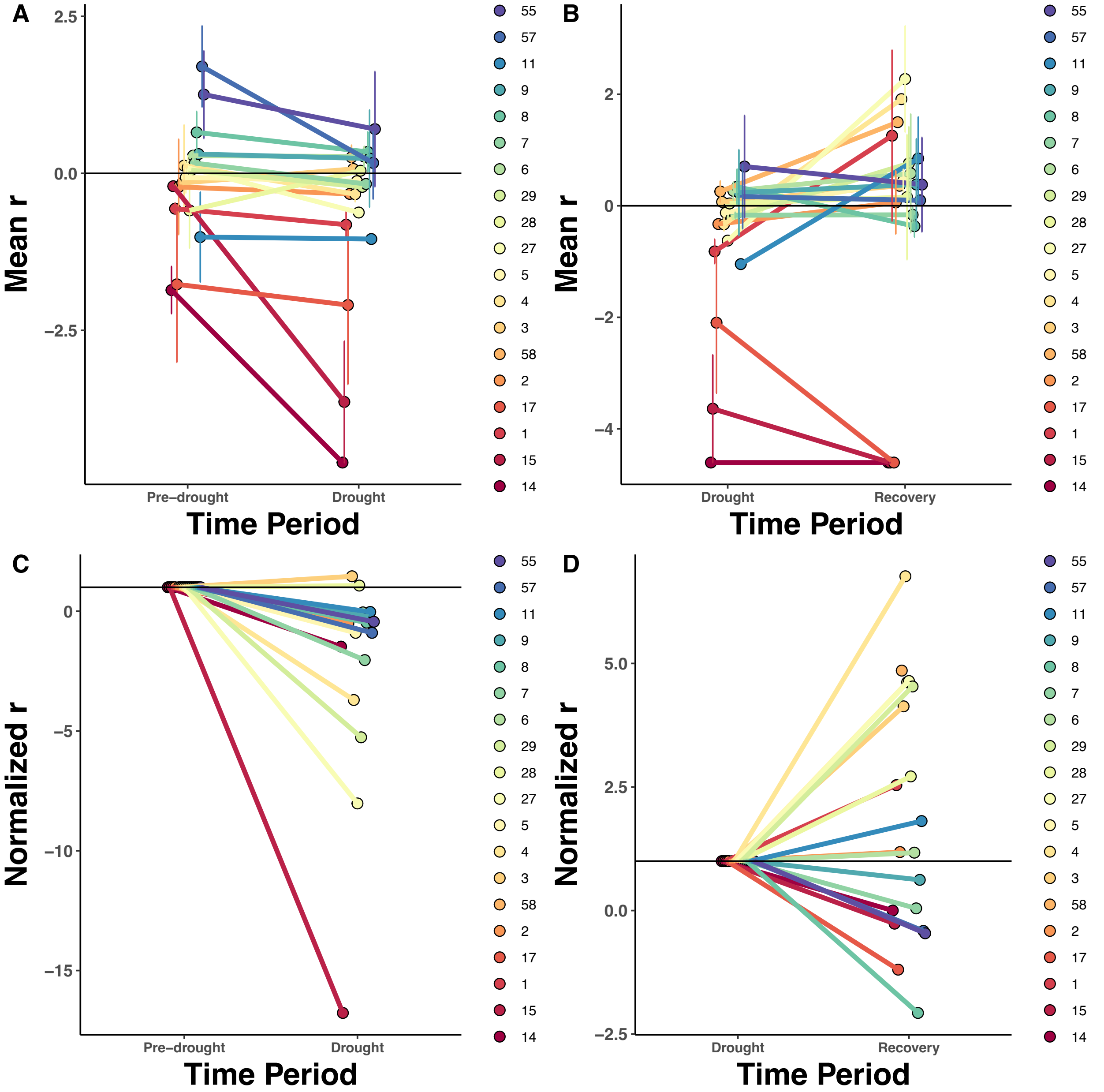

**Fig. S1.** ***Mimulus cardinalis*** **population decline and recovery from 2010-2018.** Intrinsic population growth rate (λ) for 19 populations. Colors indicate latitude of origin, from blue in the north to red in the south. Population codes are given for each population (Table S3). Demographic trends are given for (A) population decline from pre-drought (2010-2012) to drought period (2012-2015), (B) population recovery from drought to recovery period (2015-2018). Normalized r, where r at the starting period was scaled to 1, are given for (C) pre-drought to drought, and (D) drought to recovery periods. Population replacement rate (the value at which a population is projected to be stable, neither growing nor shrinking) is given as a dashed horizontal line at $r = 0$ in panels A-

B. In panels C-D, the horizontal dashed line indicates the value at which population growth does not change from starting values (normalized r=1).

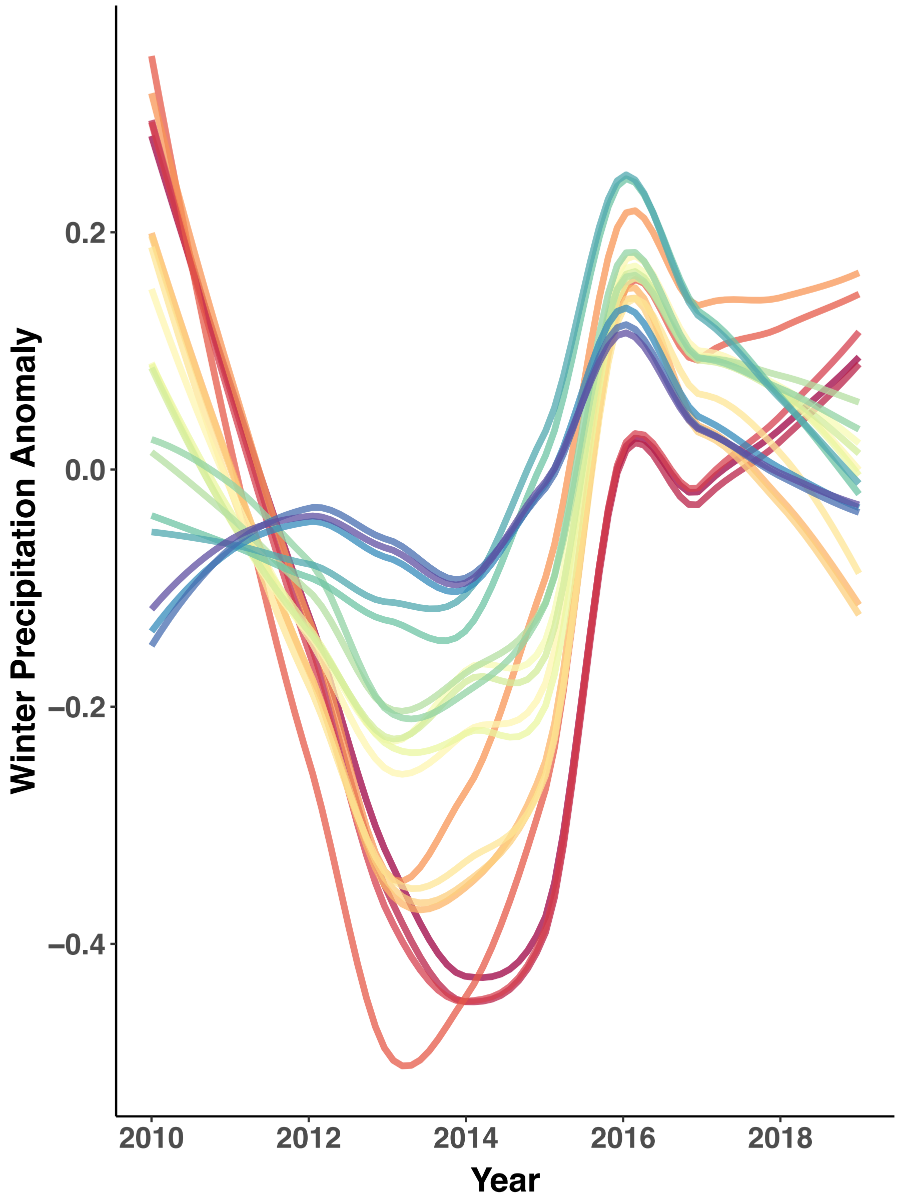

**Fig. S2** Mean annual precipitation anomaly (mm) across 19 populations of *M. cardinalis* during record-setting drought and recovery. Each curve represents the precipitation time series for one population. Rainbow colouring indicates latitudinal position (red colors = lower latitudes, blue colors = higher latitudes). See Fig. C for locations and population codes.

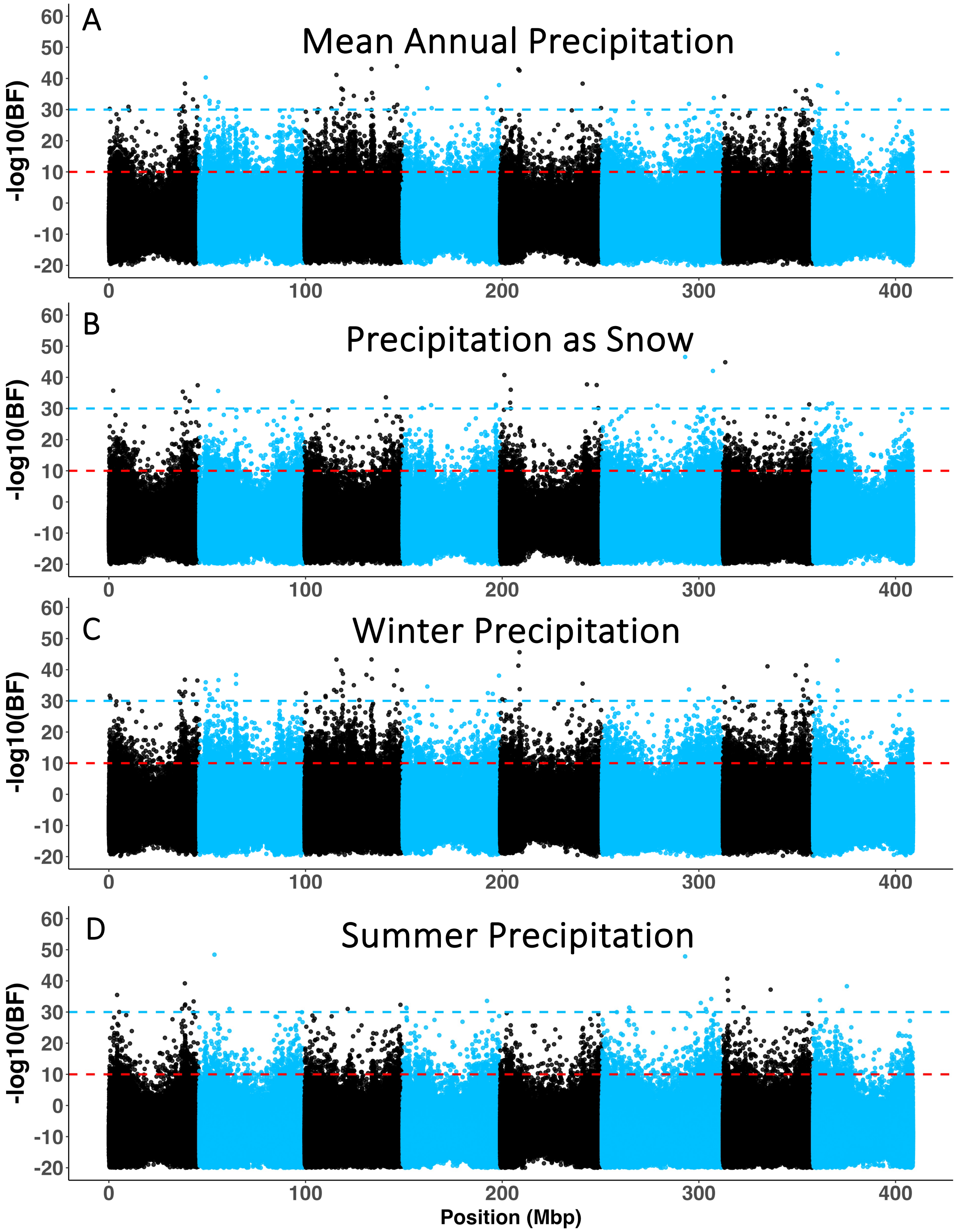
A
Mean Annual Precipitation
B
Precipitation as Snow
C
Winter Precipitation
D
Summer Precipitation
-log10(BF)
60
50
40
30
20
10
0
-10
-20
0
100
200
300
400
Position (Mbp)

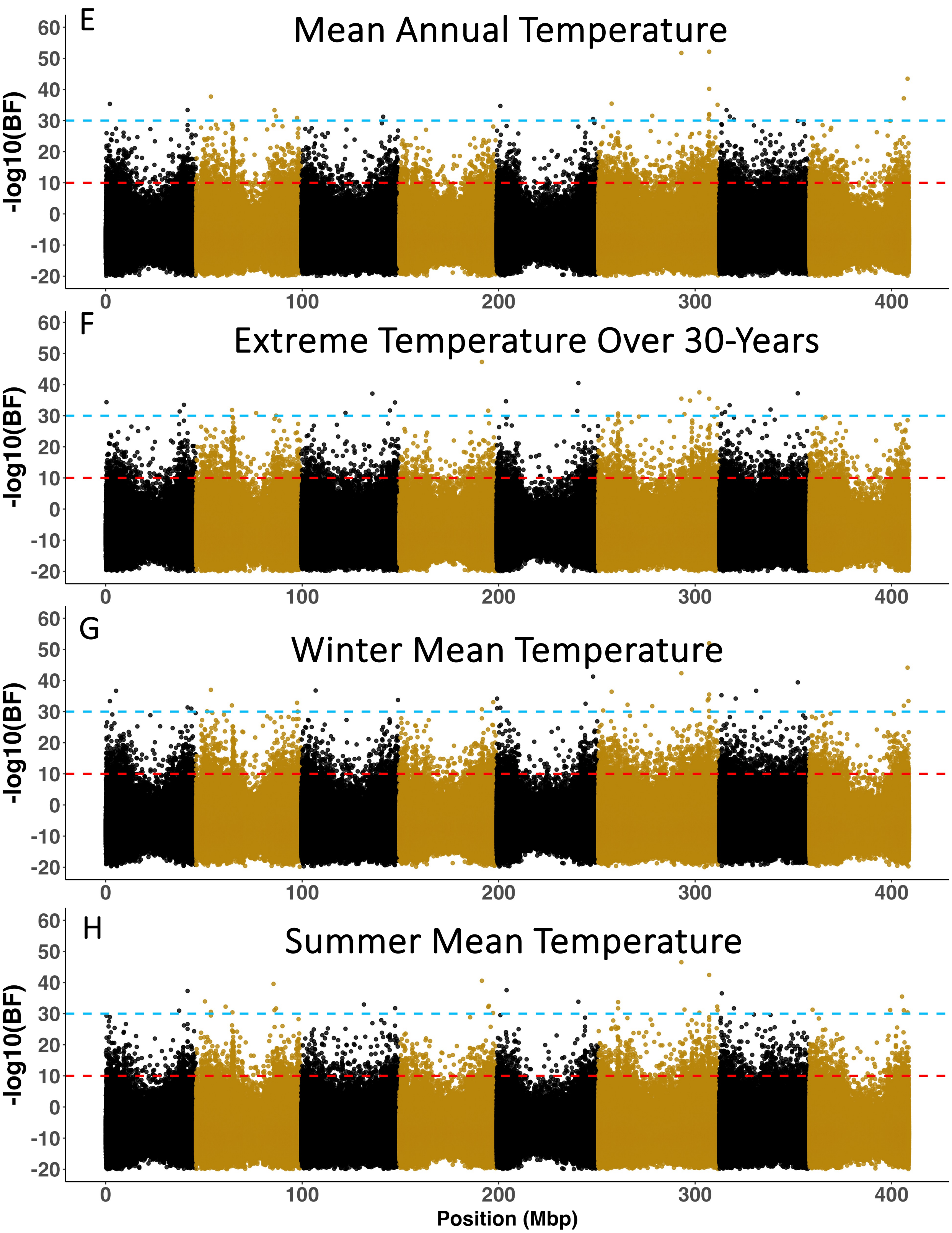
E
Mean Annual Temperature
F
Extreme Temperature Over 30-Years
G
Winter Mean Temperature
H
Summer Mean Temperature
-log10(BF)
60
50
40
30
20
10
0
-10
-20
0
100
200
300
400
Position (Mbp)

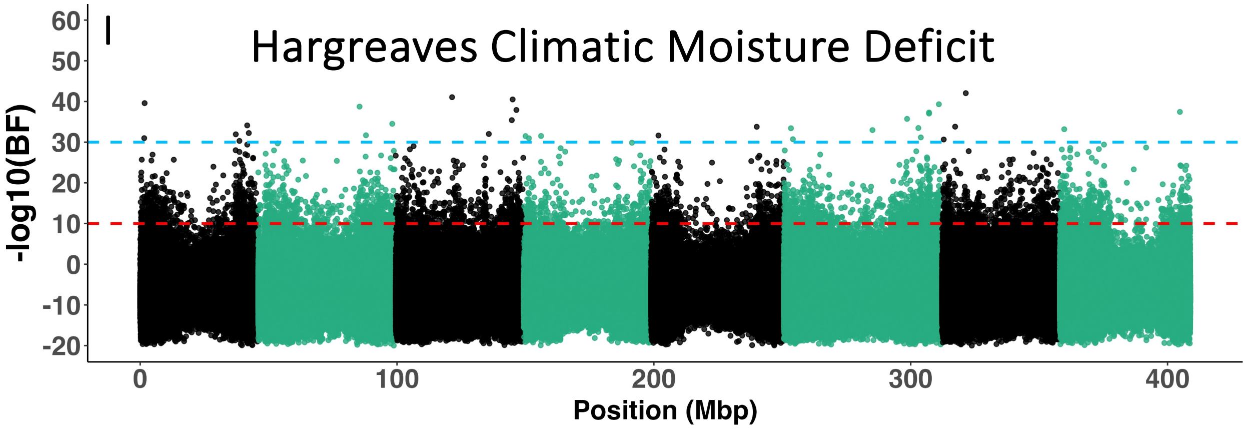

**Fig. S3. BayPass GEA Manhattan plots across nine climate variables.** -$Log_{10}$Bayes Factors (BF) given for 2,156,443 SNPs across eight chromosomes for (A) mean annual precipitation, (B) precipitation as snow, (C) winter precipitation, (D) summer precipitation, (E) mean annual temperature, (F) extreme temperature over 30 years, (G) winter mean temperature, (H) summer mean temperature, and (I) Hargreaves climatic moisture deficit. Blue dashed line = -$log_{10}$BF>30 threshold. Red dashed line = -$log_{10}$(BF)>10 threshold. Position across the genome is given in megabase pairs. SNPs across different chromosomes are coded in alternating black and colored formatting. Blue plots = SNP associations with climate variables measuring precipitation. Dark yellow plots = SNP associations with climate variables measuring temperature. Green plot = SNP associations with composite metric that involves both temperature and precipitation.

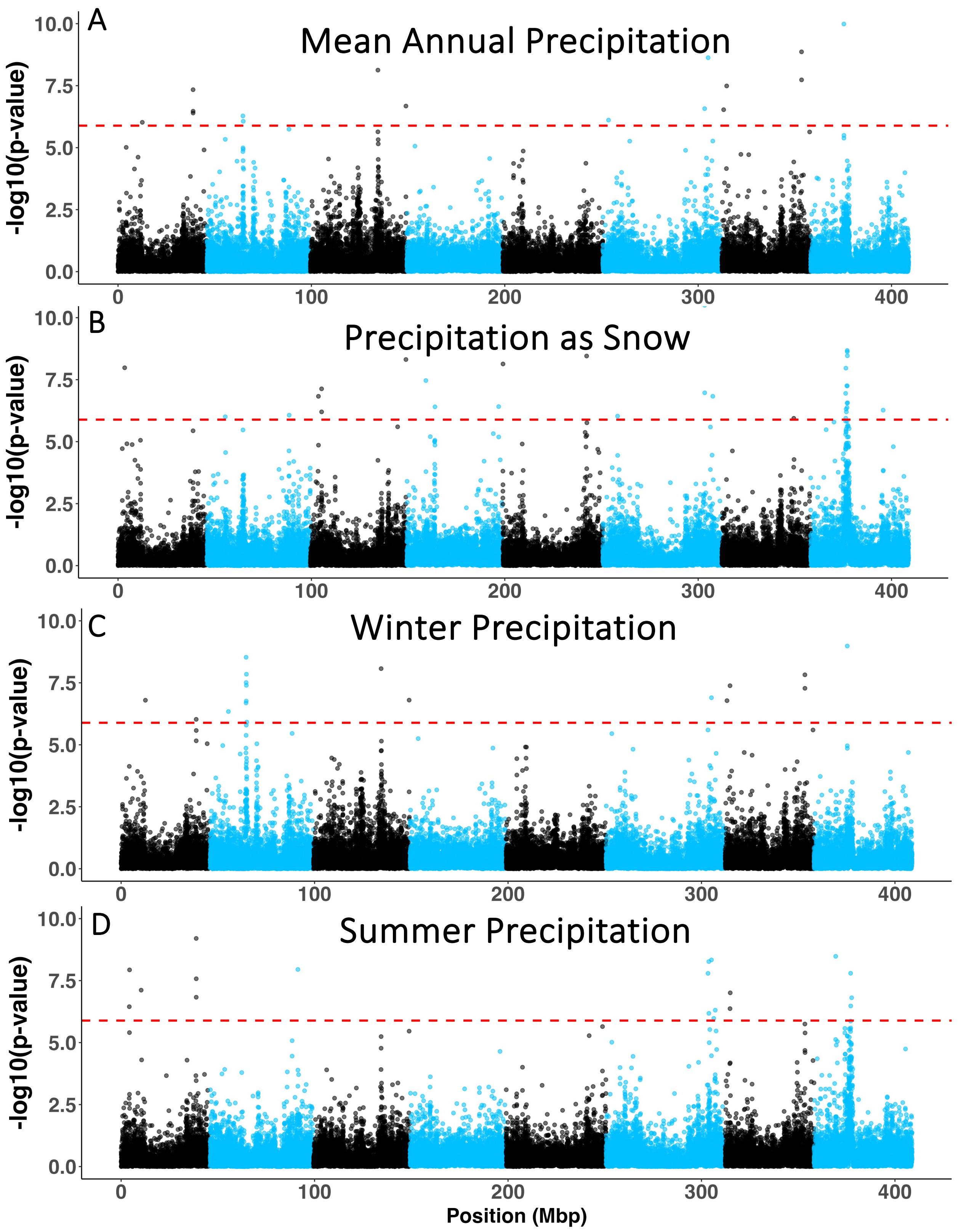

A
Mean Annual Precipitation
B
Precipitation as Snow
C
Winter Precipitation
D
Summer Precipitation
-log10(p-value)
10.0
7.5
5.0
2.5
0.0
0
100
200
300
400
Position (Mbp)

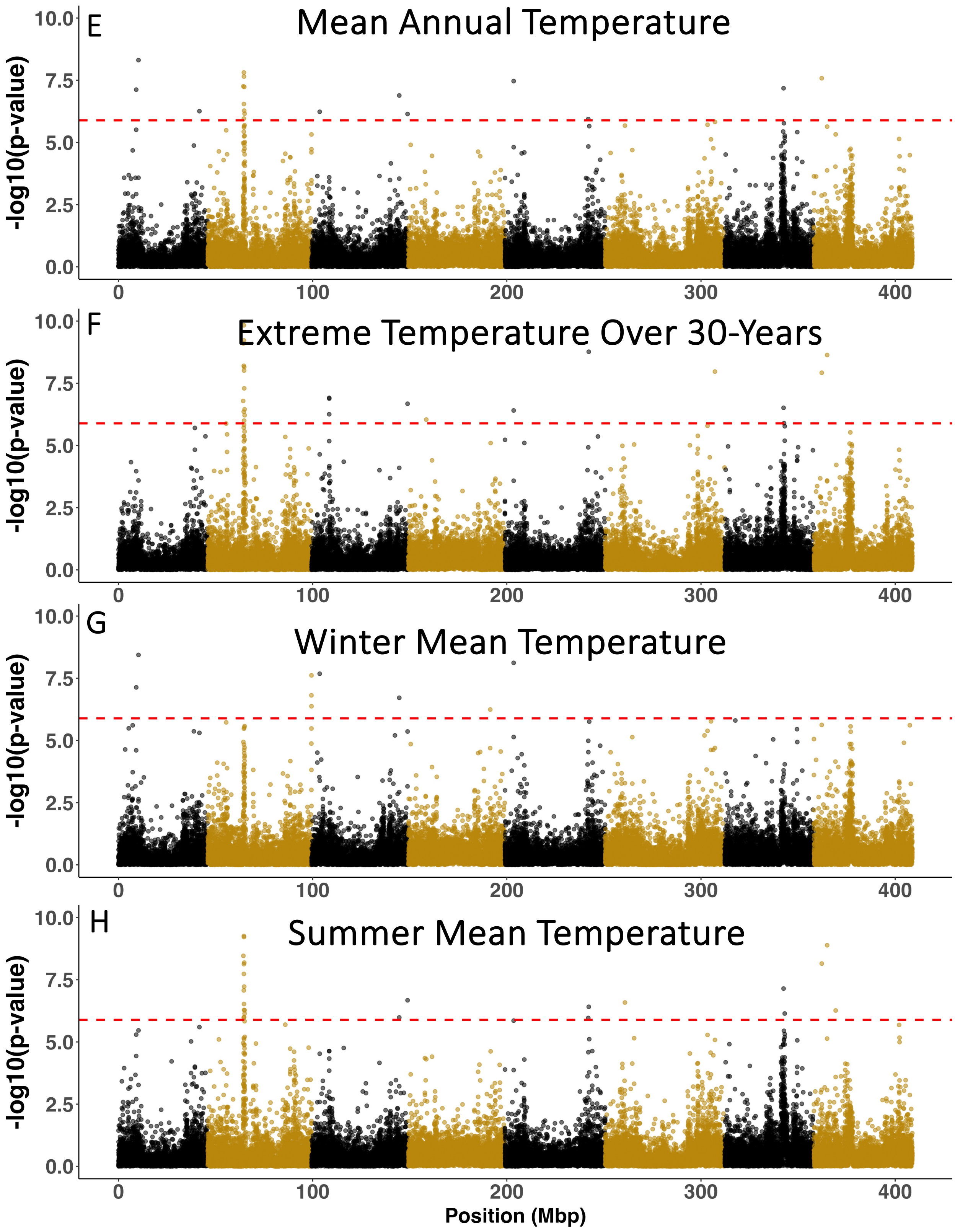
E
Mean Annual Temperature
F
Extreme Temperature Over 30-Years
G
Winter Mean Temperature
H
Summer Mean Temperature
-log10(p-value)
0.0
2.5
5.0
7.5
10.0
0
100
200
300
400
Position (Mbp)

416

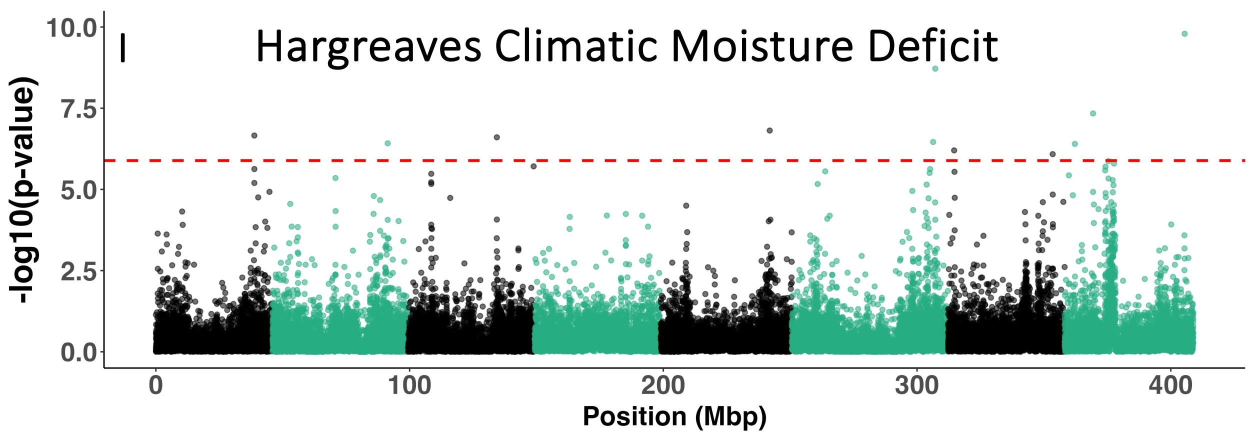

**Fig. S4. WZA genome-wide association mapping Manhattan plots across nine climate variables.** $-\text{Log}_{10}$ p-value given for 2,156,443 SNPs across eight chromosomes for (A) mean annual precipitation, (B) precipitation as snow, (C) winter precipitation, (D) summer precipitation, (E) mean annual temperature, (F) extreme temperature over 30 years, (G) winter mean temperature, (H) summer mean temperature, and (I) Hargreaves climatic moisture deficit. Red dashed line = Bonferroni significance threshold. Position across the genome is given in megabase pairs. SNPs across different chromosomes are coded in alternating black and colored formatting. Blue plots = SNP associations with climate variables measuring precipitation. Dark yellow plots = SNP associations with climate variables measuring temperature. Green plot = SNP associations with composite metric that involves both temperature and precipitation.

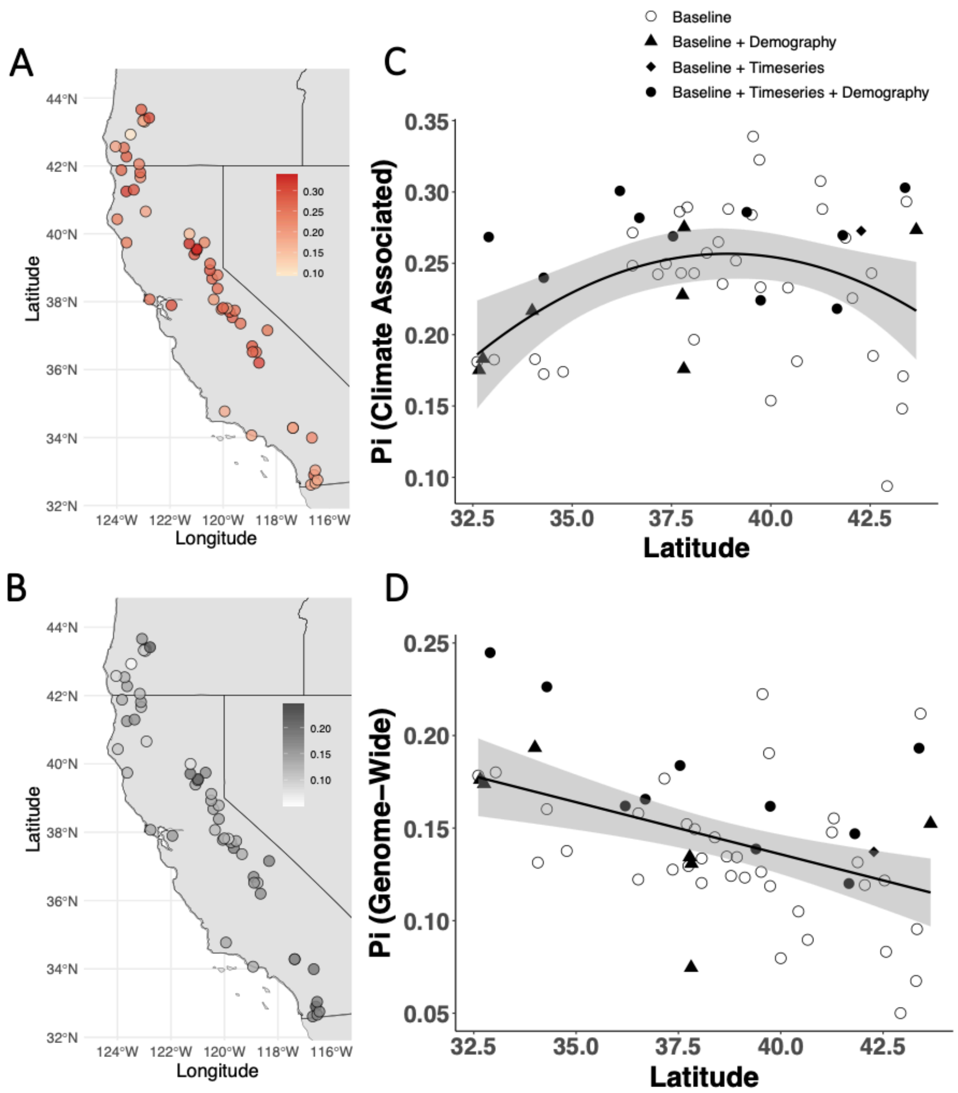

**Fig. S5. Nucleotide diversity (π) across the range of *Mimulus cardinalis* prior to exceptional drought.** π for (A) climate-associated SNPs and (B) genome-wide SNPs is given across the range with warmer colors representing larger values. π is plotted across latitude for (C) climate-associated SNPs and (D) genome-wide SNPs with the type of population indicated by legend. Baseline refers to populations that were only sequenced prior to drought. Timeseries refers to populations sequenced 2010 to 2016. Demography refers to populations for which annual growth rates were estimated before, during and after exceptional drought.

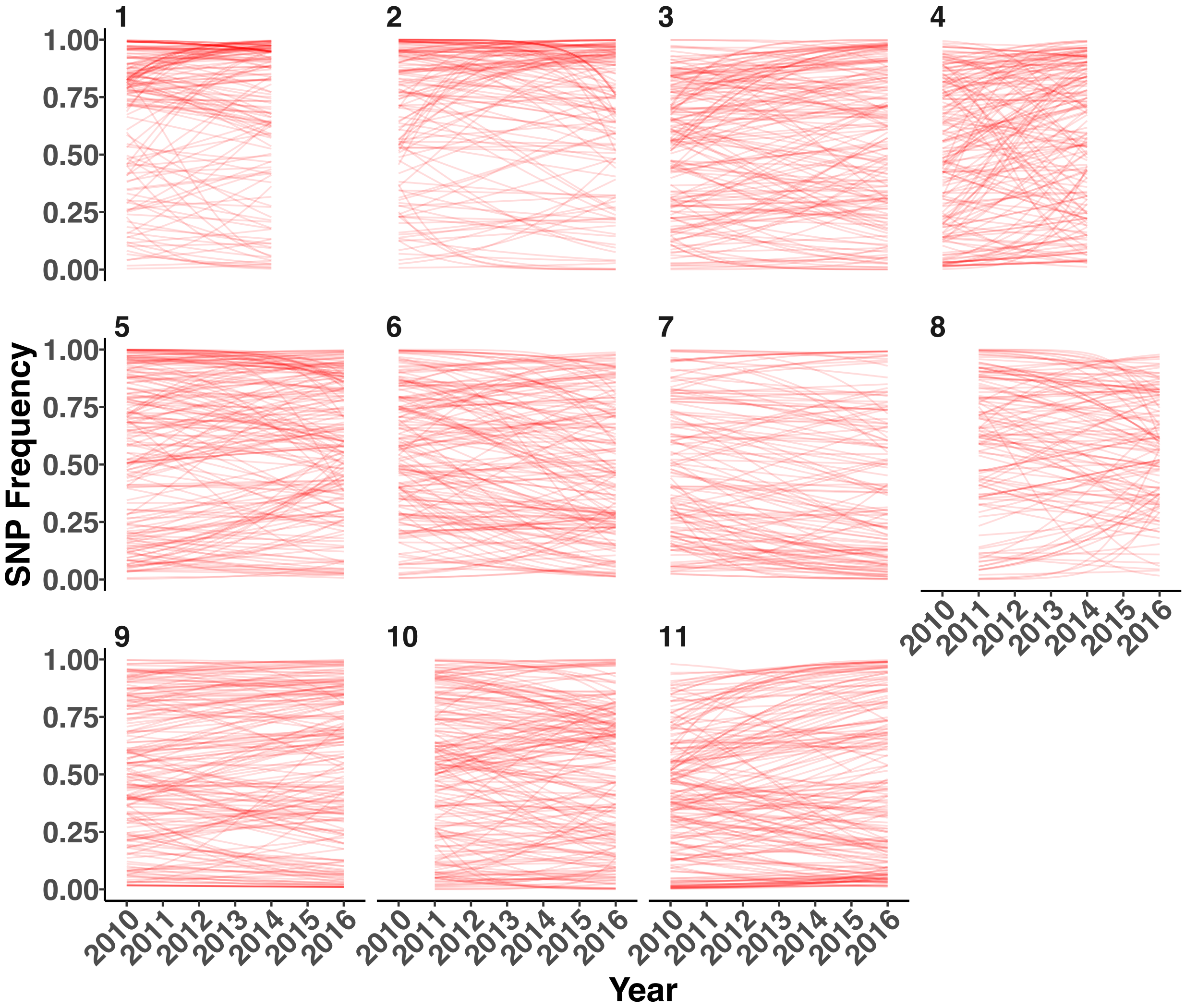

**Fig. S6. SNP frequency change through extreme drought across climate-associated SNPs.** Fitted lines from generalized linear models are shown for each of 215 climate-associated SNP frequencies over time. Each line represents a single regression of frequency vs. year for one SNP within the given population. Eleven Time Series populations are shown in ascending latitudinal order. Population codes are given for each population (Table S3). Data points are not shown.

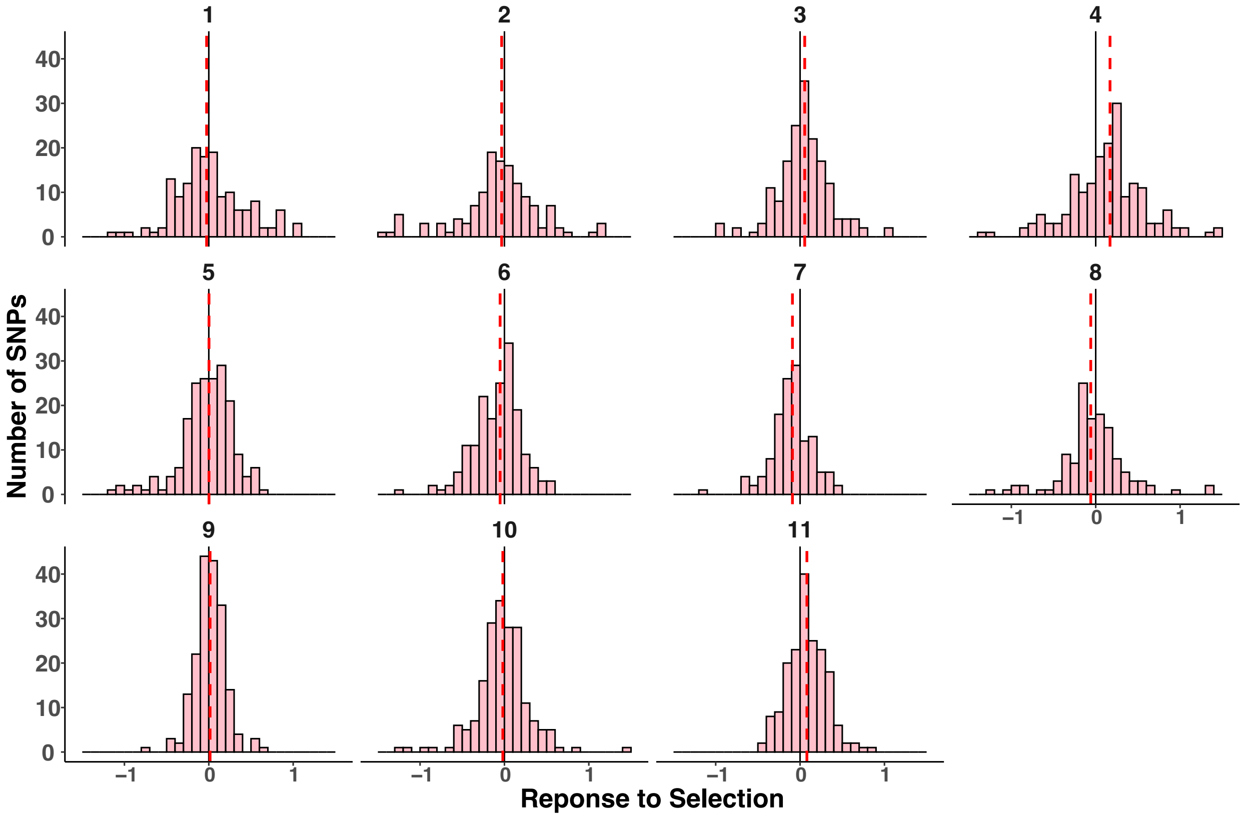

**Fig. S7. Distribution of the response to selection (S) seen during rapid drought evolution across the climate-associated SNPs.** Histograms are shown for the 11 timeseries populations in ascending latitudinal order. Light red histograms give the number of observed SNPs within 0.1 S. Positive S indicates directional selection towards the drought/heat-associated allele. Black solid line is S=0. Dashed red line is the median S across the entire distribution. Population codes are given for each population (Table S3).

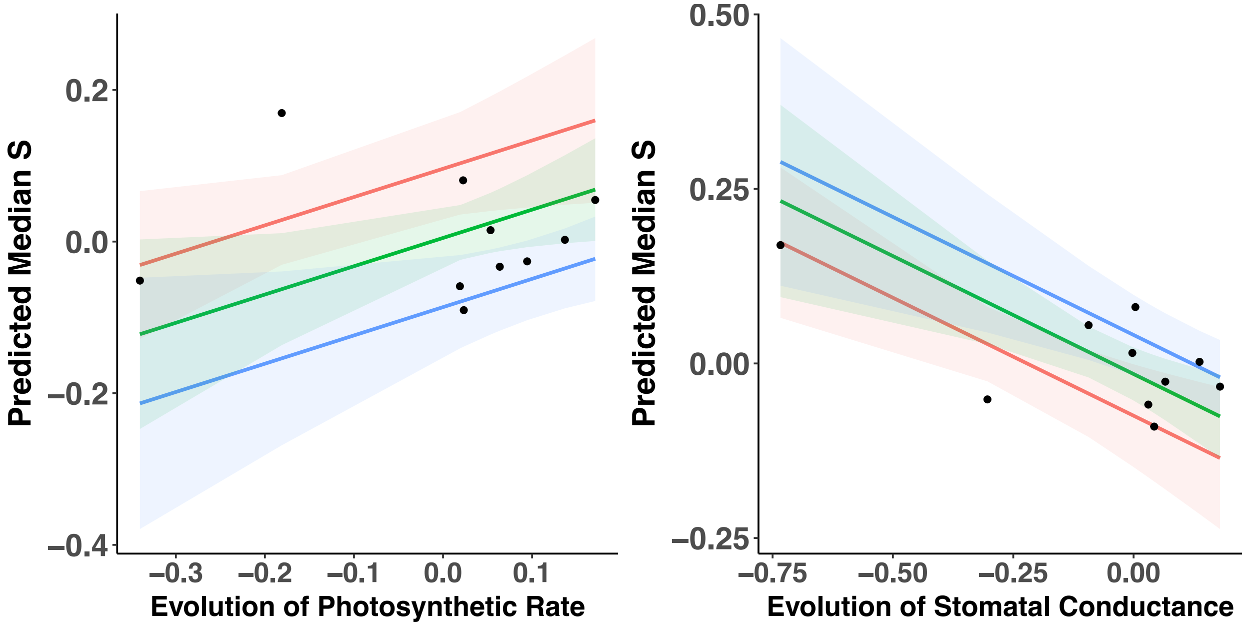

**Fig. S8 Evolution of photosynthetic rate and stomatal conductance are associated with differences in genomic response to selection (predicted median S).** Positive median S indicates evolution of increased frequency of alleles associated with drier or hotter historical conditions. Multiple regression identified two traits whose evolution is associated with median S. Evolution of each trait is quantified as the slope of trait versus year estimated from a linear mixed model fitted to data from a resurrection experiment, where lineages from each population sampled before, during and after extreme drought were phenotyped in a greenhouse common garden with a drought-simulating watering regime (Anstett et al., 2021). Positive values depict evolution of greater trait values over time (e.g., evolution towards higher photosynthetic carbon assimilation rate or higher stomatal conductance). Predictions from the model relating evolution of gas exchange traits to median S were visualized using the ggeffects package (*54*) for high (blue), medium (green) and low (red) effects of the other variable. Model predictions for each focal predictor were estimated at the mean of the non-focal predictor using the margin = "mean_reference" command.

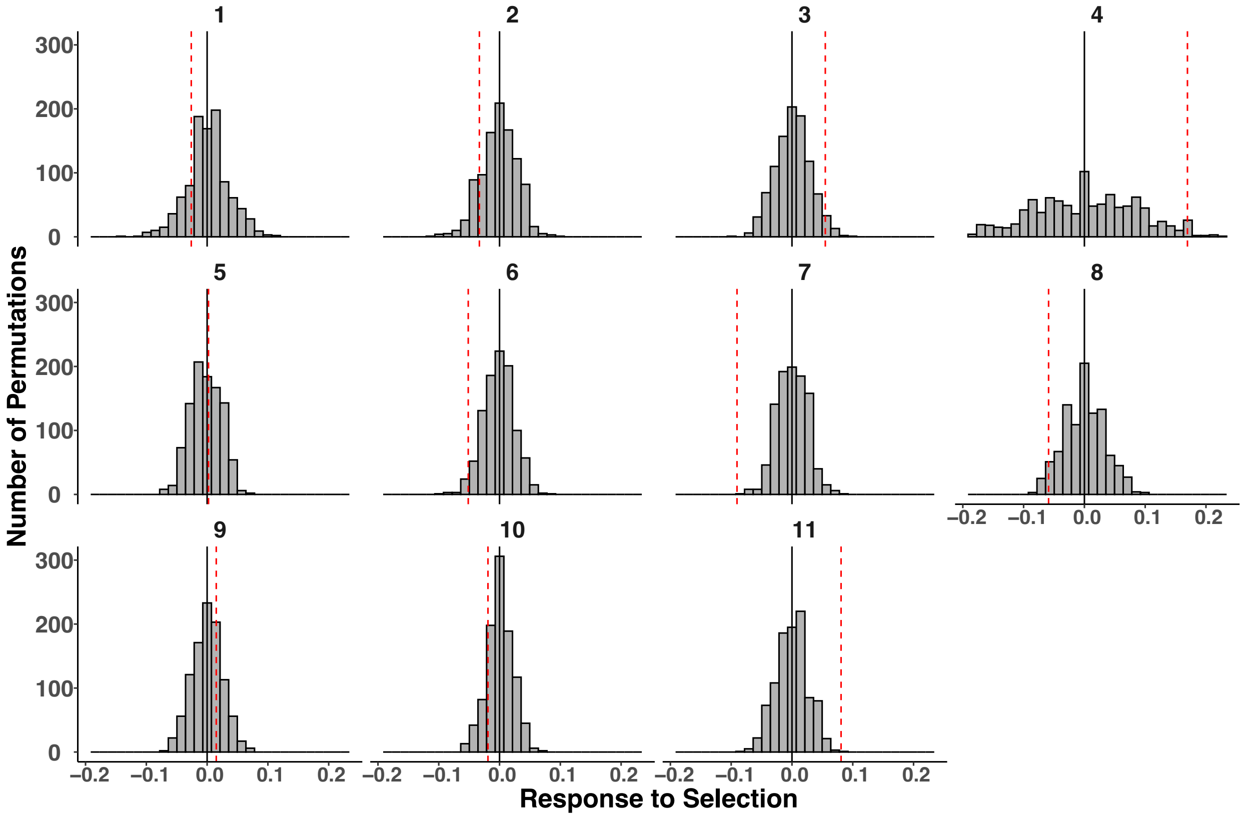

**Fig. S9. Comparison of response to selection (S) for non-climate-associated and climate-associated SNP.** The distribution of the median S of non-climate associated SNPs is given for 1000 random-stratified permutations. Red dotted vertical lines = observed mean S of the distribution of climate-associated SNPs (as in Figs. S7 and S8). Black solid line represents 0. Results are shown for the 11 Time Series populations in ascending latitudinal order. Population codes are given for each population (Table S3).

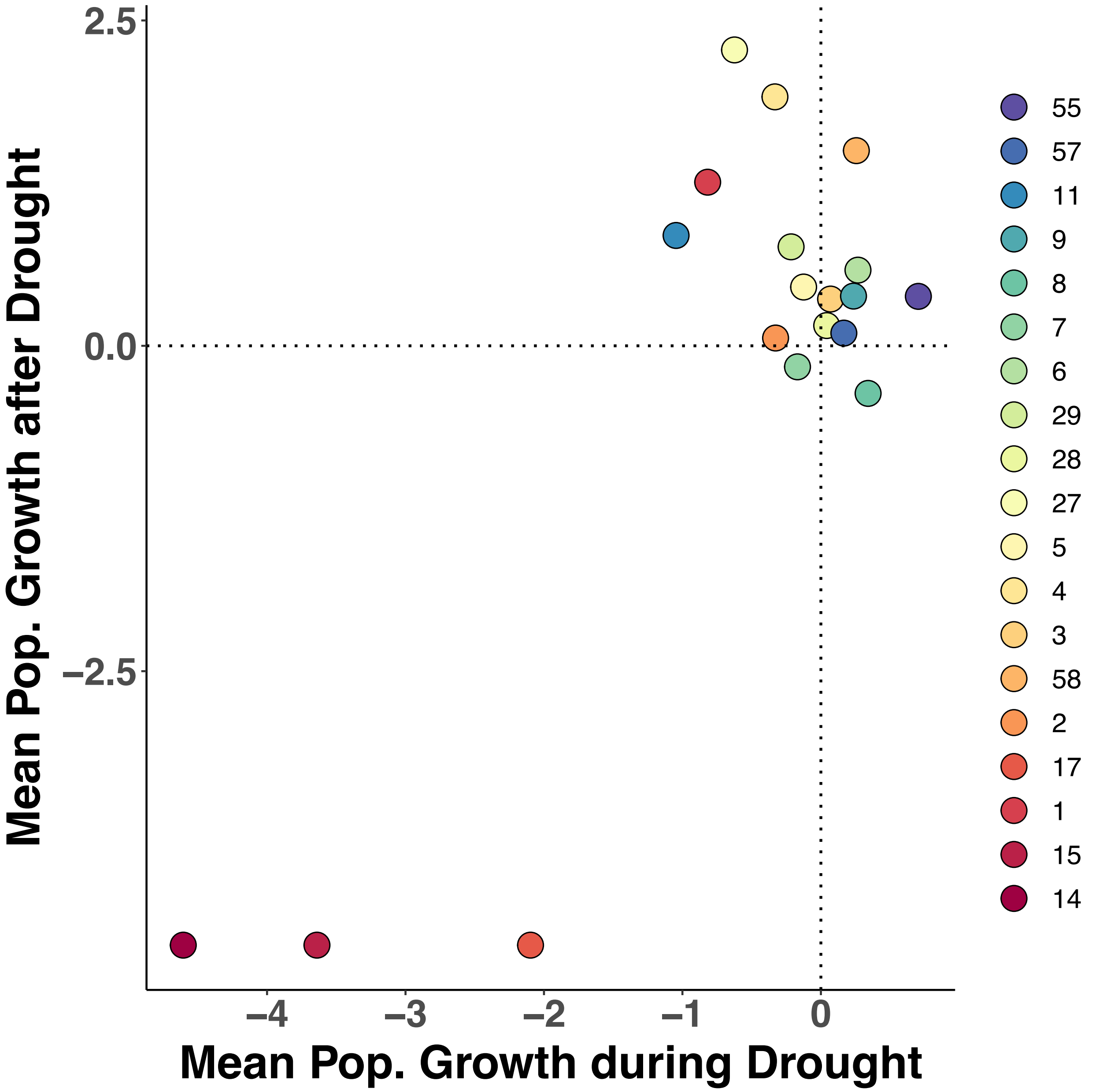

**Fig. S10. Comparison of *Mimulus cardinalis* population decline versus recovery.** Each point represents one population color coded by latitude (red = lower latitudes, blue = higher latitudes). Population codes are given for each population (Table S3). Mean population growth (*r*) for decline is given for the mean of 2012-13, 2013-14, and 2014-15 transitions. Mean *r* for recovery is given for the mean of 2015-16, 2016-17, and 2017-18 transitions.

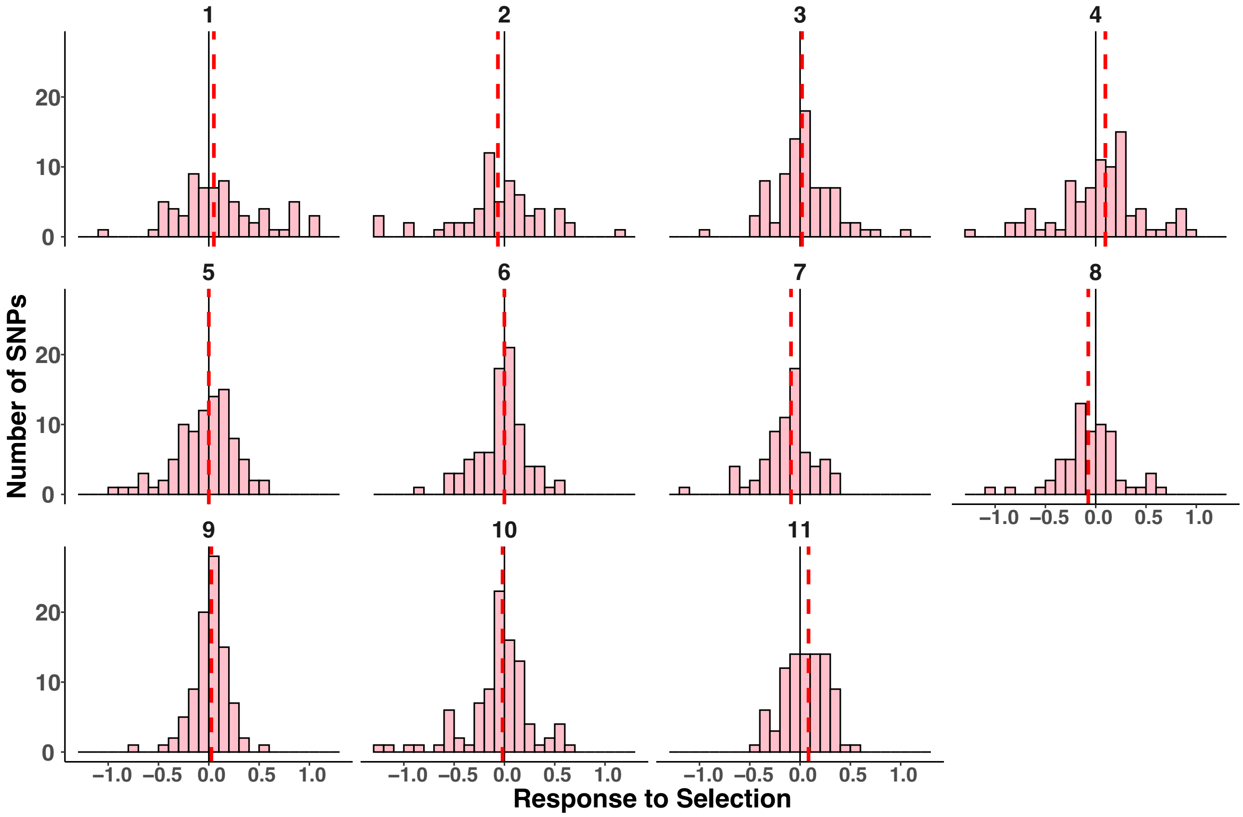

**Fig. S11. Distribution of the response to selection (S) seen during rapid drought evolution across the climate-associated SNPs, when climate-associated SNPs are identified only for the three climate variables that explain demographic decline during drought.** Results given for all SNPs monotonically changing with one or more climate variables for each of the 11 timeseries populations in ascending latitudinal order. Light red histograms give the number of observed SNPs within 0.1 S. Positive S indicates directional selection towards the drought/heat-associated allele. Black solid line is S=0. Dashed red line is the median S across the entire distribution. Population codes are given for each population (Table S3).

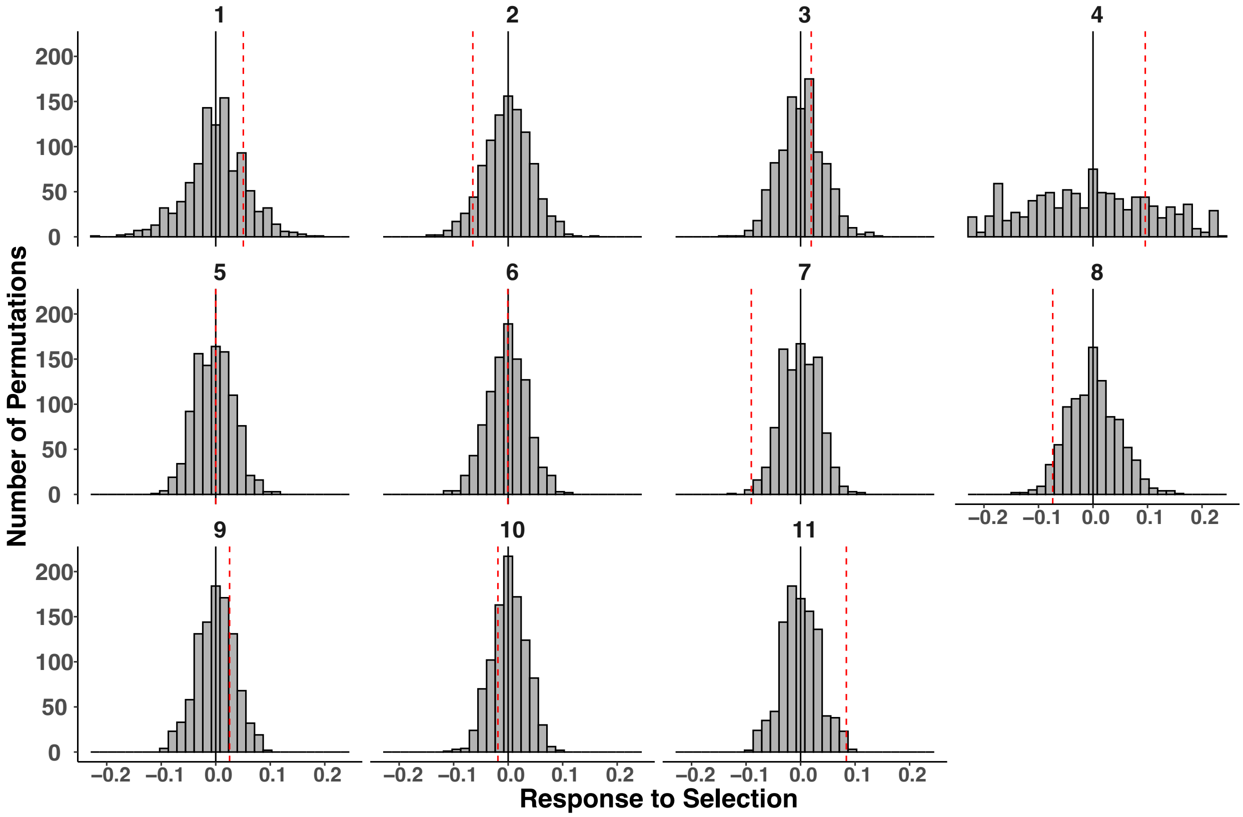

**Fig. S12. Comparison of S for non-climate-associated and climate-associated SNP, when climate-associated SNP are identified only for the three climate variables that explain demographic decline during drought.** The distribution of the median S of non-climate associated SNPs is given for 1000 random-stratified permutations. Red dotted vertical lines = observed mean S of the distribution of climate-associated SNPs (as in Figs. S7 and S8). Black solid line represents 0. Results are shown for the 11 Time Series populations in ascending latitudinal order. Population codes are given for each population (Table S3).

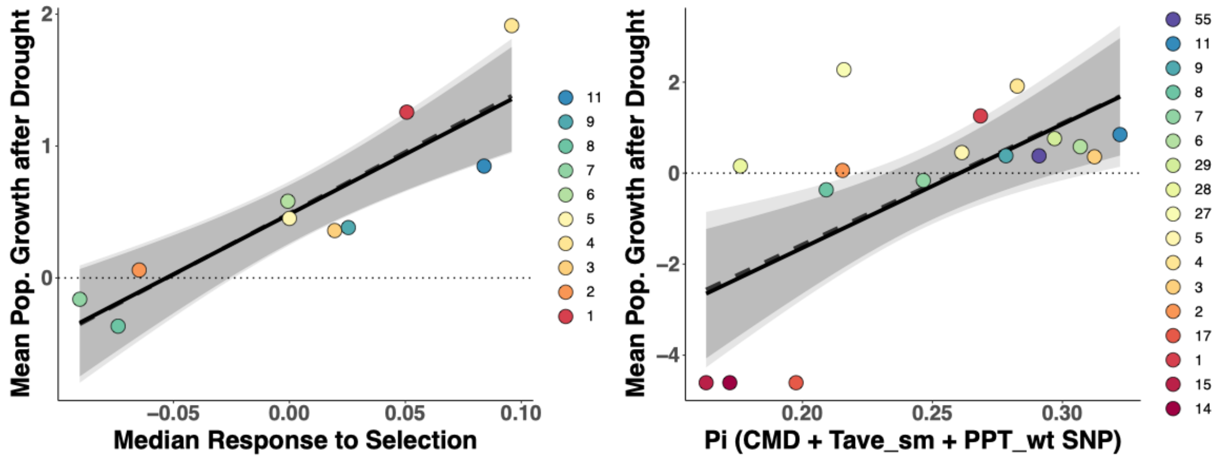

**Fig. S13. Sensitivity analysis of selection and nucleotide diversity (Pi) explaining *Mimulus cardinalis* population growth rate after drought when SNP are identified for only those climate variables that significantly predict demographic decline during drought.** Each point represents one population color coded by latitude (red = lower latitudes, blue = higher latitudes). Population codes are given for each population (Table S3). Mean *r* for recovery is given for the mean of 2015-16, 2016-17, and 2017-18 transitions. Solid lines and darker shading depict predicted the slope and its 95% confidence intervals from robust regression. Dashed line and lighter shading depicts the slope and its 95% confidence intervals from ordinary least squares regression. The dashed line depicts rate of replacement ($r$=0).

**Table S1. Means for intrinsic rate of population growth (*r*) before drought (2010-11 and 2011-12 projection intervals), during drought (2012-13, 2013-14, 2014-15 projection intervals) and recovery (2015-16, 2016-17, 2017-18 projection intervals), and mean generation time (estimated as the expected age at reproduction for a cohort), for each population. Populations are listed in order of ascending latitude.**

| Population | Population ID | Latitude (°N) | Longitude (°W) | Mean *r* Pre-drought | Mean *r* Drought | Mean *r* Recovery | Mean gen. time |
|---|---|---|---|---|---|---|---|
| Hauser Creek | 14 | 32.6582 | -116.5323 | -1.86 | -4.61 | -4.61 | 2.02 |
| Kitchen Creek | 15 | 32.7521 | -116.4522 | -0.20 | -3.64 | -4.61 | 2.50 |
| Sweetwater River | 1 | 32.8991 | -116.5865 | -0.56 | -0.82 | 1.26 | 2.52 |
| Whitewater Canyon | 17 | 33.9933 | -116.6627 | -1.77 | -2.10 | -4.61 | 2.55 |
| West Fork Mojave River | 2 | 34.2843 | -117.3754 | -0.22 | -0.33 | 0.06 | 2.80 |
| South Fork Middle Fork Tule | 58 | 36.1376 | -118.5741 | NA | 0.26 | 1.50 | 4.59 |
| North Fork Middle Fork Tule | 3 | 36.2008 | -118.6509 | -0.15 | 0.07 | 0.36 | 3.69 |
| Redwood Creek | 4 | 36.6895 | -118.9096 | 0.12 | -0.33 | 1.91 | 2.75 |
| Wawona | 5 | 37.5382 | -119.6514 | -0.07 | -0.12 | 0.45 | 3.25 |
| Buck Meadows | 27 | 37.7772 | -120.0633 | 0.09 | -0.62 | 2.27 | 3.33 |
| Carlon | 28 | 37.8101 | -119.8568 | -0.59 | 0.04 | 0.16 | 2.63 |
| Rainbow Pool | 29 | 37.8188 | -120.0073 | 0.05 | -0.22 | 0.76 | 3.04 |
| Oregon Creek | 6 | 39.3970 | -121.0820 | 0.28 | 0.27 | 0.58 | 4.18 |
| Little Jamison Creek | 7 | 39.7430 | -120.7040 | 0.16 | -0.17 | -0.16 | 5.14 |
| Deep Creek | 8 | 41.6655 | -123.1134 | 0.65 | 0.34 | -0.37 | 4.34 |
| O'Neil Creek | 9 | 41.8098 | -123.1189 | 0.31 | 0.24 | 0.38 | 4.61 |
| Rock Creek | 11 | 43.3788 | -122.9521 | -1.01 | -1.05 | 0.85 | 2.43 |
| Canton Creek | 57 | 43.4137 | -122.7804 | 1.70 | 0.17 | 0.10 | 4.11 |
| Coast Fork of Willamette | 55 | 43.6517 | -123.0880 | 1.25 | 0.70 | 0.38 | 2.74 |

**Table S2. Association between climate anomalies and *M. cardinalis* population demography. Associations were estimated for 19 populations with demographic observations using ordinary least squares (OLS) regression and robust regression, which minimizes the influence of potential outliers. Climate anomalies were calculated as the difference between values during the observation period and historical mean values from 1979 – 2010.**

| | Drought Period | | | | | Recovery Period | | | | |
|---|---|---|---|---|---|---|---|---|---|---|
| **Climate Anomaly** | **OLS Slope** | **OLS P-value** | **OLS adj-$R^2$** | **Robust Slope** | **Robust P-value** | **OLS Slope** | **OLS P-value** | **OLS adj-$R^2$** | **Robust Slope** | **Robust P-value** |
| Mean Annual Temperature | 0.23 | 0.361 | -0.01 | 0.08 | 0.559 | 0.02 | 0.944 | -0.06 | 0.01 | 0.944 |
| Mean Annual Precipitation | 0.29 | 0.238 | 0.03 | 0.19 | 0.132 | **0.50** | **0.037** | **0.20** | 0.38 | 0.137 |
| Precipitation as Snow | -0.18 | 0.475 | -0.03 | -0.05 | 0.743 | 0.17 | 0.490 | -0.03 | -0.04 | 0.704 |
| Hargreaves Climatic Moisture Deficit | -0.47 | 0.051 | 0.17 | -0.29 | 0.097 | -0.18 | 0.482 | -0.03 | -0.03 | 0.780 |
| Winter Mean Temperature | 0.01 | 0.959 | -0.06 | -0.04 | 0.757 | -0.08 | 0.743 | -0.06 | 0.00 | 0.983 |
| Summer Mean Temperature | **0.61** | **0.007** | **0.33** | **0.35** | **0.027** | 0.43 | 0.077 | 0.13 | 0.15 | 0.405 |
| Winter Precipitation | **0.56** | **0.015** | **0.28** | **0.43** | **0.001** | 0.45 | 0.062 | 0.15 | 0.13 | 0.431 |
| Summer Precipitation | -0.08 | 0.749 | -0.06 | -0.06 | 0.678 | -0.22 | 0.379 | -0.01 | -0.07 | 0.489 |

p<0.05 is given in bold. p<0.10 is underlined.

**Table S3. Population representation across demography, baseline, and timeseries datasets.**

| Population | Population ID | Latitude | Longitude | Demography | Baseline Genomes | Collection Year | Timeseries | Selection Comparison to Demographic Recovery | Baseline Pi Comparison to Demography Recovery |
|---|---|---|---|---|---|---|---|---|---|
| Sweetwater River | 1 | 32.89928 | -116.5849 | Yes | 9 | 2010 | Yes | Yes | Yes |
| West Fork Mojave River | 2 | 34.28425 | -117.37539 | Yes | 5 | 2010 | Yes | Yes | Yes |
| North Fork Middle Fork Tule River | 3 | 36.20081 | -118.65092 | Yes | 7 | 2010 | Yes | Yes | Yes |
| Redwood Creek | 4 | 36.69096 | -118.90961 | Yes | 5 | 2010 | Yes | Yes | Yes |
| Wawona | 5 | 37.539 | -119.654 | Yes | 7 | 2010 | Yes | Yes | Yes |
| Oregon Creek | 6 | 39.39442 | -121.08302 | Yes | 10 | 2010 | Yes | Yes | Yes |
| Little Jamison Creek | 7 | 39.74298 | -120.70401 | Yes | 8 | 2010 | Yes | Yes | Yes |
| Deep Creek | 8 | 41.66546 | -123.11341 | Yes | 4 | 2011 | Yes | Yes | Yes |
| O'Neil Creek | 9 | 41.80979 | -123.11887 | Yes | 9 | 2010 | Yes | Yes | Yes |
| Deer Creek | 10 | 42.27411 | -123.63617 | Yes | 10 | 2011 | Yes | No | Yes |
| Rock Creek | 11 | 43.37876 | -122.95207 | Yes | 10 | 2010 | Yes | Yes | Yes |
| Cottonwood Creek | 13 | 32.60831 | -116.70098 | No | 11 | 2010 | No | No | No |
| Hauser Creek | 14 | 32.65822 | -116.53235 | Yes | 10 | 2010 | No | No | Yes |
| Kitchen Creek | 15 | 32.75206 | -116.45221 | Yes | 10 | 2010 | No | No | Yes |
| Chariot Canyon | 16 | 33.03596 | -116.53623 | No | 7 | 2010 | No | No | No |
| Whitewater Canyon | 17 | 33.99329 | -116.66267 | Yes | 9 | 2010 | No | No | Yes |
| Arroyo Sequit | 18 | 34.06511 | -118.93279 | No | 5 | 2010 | No | No | No |
| Tributary to West Fork Mojave River | 19 | 34.28497 | -117.37862 | No | 4 | 2010 | No | No | No |
| Manzana Creek | 20 | 34.77171 | -119.94363 | No | 2 | 2010 | No | No | No |
| Paradise Creek | 21 | 36.51776 | -118.75877 | No | 5 | 2010 | No | No | No |
| North Fork Seep | 22 | 36.52105 | -118.89359 | No | 4 | 2008 | No | No | No |
| Baker Creek | 23 | 37.15761 | -118.3334 | No | 6 | 2010 | No | No | No |

| | | | | | | | | | |
|---|---|---|---|---|---|---|---|---|---|
| Unknown Creek | 24 | 37.3592 | -119.34498 | No | 10 | 2009 | No | No | No |
| Crane Creek | 25 | 37.70377 | -119.75363 | No | 10 | 2010 | No | No | No |
| Tenaya Creek | 26 | 37.743 | -119.562 | No | 4 | 2008 | No | No | No |
| Buck Meadows | 27 | 37.7769 | -120.063 | Yes | 2 | 2010 | No | No | Yes |
| Carlon | 28 | 37.810067 | -119.8568 | Yes | 1 | 2010 | No | No | Yes |
| Rainbow Pool | 29 | 37.8188 | -120.00743 | Yes | 4 | 2009 | No | No | Yes |
| Deer Flat Creek | 30 | 37.89761 | -121.94553 | No | 2 | 2010 | No | No | No |
| Fivemile Creek | 31 | 38.06638 | -120.35368 | No | 1 | 2010 | No | No | No |
| Nicasio Creek | 32 | 38.0695 | -122.76317 | No | 7 | 2009 | No | No | No |
| North Fork Stanislaus River | 33 | 38.3872 | -120.21049 | No | 10 | 2010 | No | No | No |
| Camp Creek | 34 | 38.67988 | -120.41679 | No | 10 | 2009 | No | No | No |
| Wrights Road | 35 | 38.7854 | -120.21366 | No | 8 | 2009 | No | No | No |
| Creek to Rubicon River | 36 | 38.92881 | -120.49947 | No | 9 | 2009 | No | No | No |
| Duncan Creek | 37 | 39.12205 | -120.49242 | No | 10 | 2009 | No | No | No |
| Fiddle Creek | 38 | 39.5204 | -120.99815 | No | 2 | 2010 | No | No | No |
| Cherokee Creek | 39 | 39.55384 | -120.98785 | No | 10 | 2010 | No | No | No |
| Little North Fork Middle Fork Feather River | 40 | 39.71209 | -121.27485 | No | 5 | 2010 | No | No | No |
| South Fork Eel River | 41 | 39.74001 | -123.63228 | No | 4 | 2009 | No | No | No |
| North Feather River (Chip's Creek) | 42 | 39.99982 | -121.26988 | No | 10 | 2009 | No | No | No |
| Bear Creek | 43 | 40.43141 | -123.98378 | No | 5 | 2009 | No | No | No |
| Trinity River | 44 | 40.65827 | -122.91334 | No | 2 | 2008 | No | No | No |
| Slate Creek | 45 | 41.25024 | -123.64343 | No | 4 | 2008 | No | No | No |
| Salmon River 2 | 46 | 41.2969 | -123.36034 | No | 8 | 2009 | No | No | No |
| Smith River | 47 | 41.87876 | -123.82774 | No | 5 | 2009 | No | No | No |
| Carberry Creek | 48 | 42.053833 | -123.1629 | No | 7 | 2009 | No | No | No |
| North Fork Silver Creek | 49 | 42.53529 | -123.73016 | No | 10 | 2010 | No | No | No |
| Shasta Costa Creek | 50 | 42.57281 | -124.04742 | No | 3 | 2008 | No | No | No |

| | | | | | | | | | |
|---|---|---|---|---|---|---|---|---|---|
| Cow Creek | 51 | 42.92653 | -123.48479 | No | 1 | 2008 | No | No | No |
| Honey Creek | 52 | 43.305817 | -122.95505 | No | 2 | 2007 | No | No | No |
| North Fork Umpqua River | 53 | 43.3282 | -123.0154 | No | 3 | 2007 | No | No | No |
| Susan Creek | 54 | 43.4137 | -122.78043 | No | 1 | 2010 | No | No | No |
| Coast Fork Willamette River | 55 | 43.66 | -123.07997 | Yes | 10 | 2009 | No | No | Yes |
| Canton Creek | 57 | 43.4137 | -122.7804 | Yes | 0 | NA | No | No | No |
| South Fork Middle Fork Tule River | 58 | 36.13756 | -118.5741 | Yes | 0 | NA | No | No | No |

Number of baseline genomes and year of collection of baseline data is included for each population. Name and population ID is given for each population. "Yes" under demography indicates population has demography data collected from 2010 to 2019. "Yes" under timeseries indicates that the population has genomes resequenced from 2010 to 2016 (see Table S3).

**Table S4. Number of SNPs retained per climate association after SNP selection protocols.**

| Environmental Variable | Monotonic SNPs retained (for Response to Selection) | SNPs retained (For PI Calculation) |
|---|---|---|
| Mean annual temperature | 18 | 36 |
| Mean annual precipitation | 49 | 74 |
| Precipitation as snow | 31 | 48 |
| Extreme temperature | 22 | 41 |
| Hargreaves climatic moisture deficit | 29 | 47 |
| Winter mean temperature | 28 | 45 |
| Summer mean temperature | 22 | 41 |
| Winter precipitation | 54 | 78 |
| Summer precipitation | 28 | 50 |
| **Total Unique** | **215** | **341** |

**Table S5. Number of genomes sequenced from 2010 to 2016 across the 11 timeseries populations.**

| Population | Population ID | 2010 | 2011 | 2012 | 2013 | 2014 | 2015 | 2016 |
|---|---|---|---|---|---|---|---|---|
| Sweetwater River | 1 | 9 | 7 | 6 | 6 | 10 | NA | NA |
| West Fork Mojave River | 2 | 5 | 6 | NA | NA | 9 | 7 | 3 |
| North Fork Middle Fork Tule River | 3 | 7 | 2 | 3 | 5 | 8 | 9 | 4 |
| Redwood Creek | 4 | 5 | 2 | 9 | 4 | 1 | NA | NA |
| Wawona | 5 | 7 | NA | 10 | NA | 6 | 6 | 6 |
| Oregon Creek | 6 | 10 | NA | NA | 2 | 8 | 2 | 1 |
| Little Jamison Creek | 7 | 8 | NA | 8 | 10 | 9 | NA | 9 |
| Deep Creek | 8 | NA | 4 | NA | NA | 1 | 5 | 9 |
| O'Neil Creek | 9 | 8 | NA | 4 | 4 | 5 | 7 | 10 |
| Deer Creek | 10 | NA | 10 | 10 | NA | 8 | 9 | 9 |
| Rock Creek | 11 | 10 | NA | 2 | NA | 5 | 10 | 10 |

**Table S6. Statistics for median response to selection (S) on climate-associated alleles across the 11 timeseries populations.**

| Population | Population ID | Latitude | Longitude | Median S | Wilcoxon P-value | Median Percentile | Median P-value |
|---|---|---|---|---|---|---|---|
| Sweetwater River | 1 | 32.89913 | −116.5865 | -0.03 | 0.97 | 0.174 | 0.826 |
| West Fork Mojave River | 2 | 34.28425 | −117.3754 | -0.03 | 0.24 | 0.142 | 0.858 |
| NFMF Tule River | 3 | 36.20081 | −118.6509 | 0.05 | **0.03** | 0.968 | **0.032** |
| Redwood Creek | 4 | 36.68952 | −118.9096 | 0.17 | **<0.001** | 0.977 | **0.023** |
| Wawona | 5 | 37.53822 | −119.6514 | 0.002 | 0.87 | 0.576 | 0.424 |
| Oregon Creek | 6 | 39.39704 | −121.082 | -0.05 | **<0.001** | **0.027** | 0.973 |
| Little Jamison Creek | 7 | 39.74298 | −120.704 | -0.09 | **<0.001** | **0** | 1 |
| Deep Creek | 8 | 41.66546 | −123.1134 | -0.06 | 0.18 | **0.039** | 0.961 |
| O'Neil Creek | 9 | 41.80979 | −123.1189 | 0.02 | 0.30 | 0.726 | 0.274 |
| Deer Creek | 10 | 42.27411 | −123.63617 | -0.02 | 0.30 | 0.16 | 0.84 |
| Rock Creek | 11 | 43.37876 | −122.9521 | 0.08 | **<0.001** | 0.999 | **0.001** |

Median S is the median climate-associated response to selection, derived from logistic regression of allele frequency across years. Wilcoxon p-value is given for the test that the median S of climate-associated loci is different from 0. Sets of non-climate associated SNPs were randomly selected 1000 times to generate 1000 null distributions of S, and the median was calculated for each of these distributions, generating a null distribution of S. Median percentile gives the percentile of the null distribution where the observed climate-associated median lies. $p<0.05$ is given in bold for evidence that the climate-associated median S is less than would be expected by drift alone. Empirical p-value is given for the test that the climate associated median is greater than the non-climate associated distribution of medians. $p<0.05$ is given in bold.

**Table S7. Statistics for multiple regression of phenotypic variables explaining median response to selection.**

| Regression | $R^2$ | F | dfn | dfd | P-value | Predictor | Coeff | SE | t | P-value |
|---|---|---|---|---|---|---|---|---|---|---|
| Anatomy | -0.23 | 0.17 | 2 | 7 | 0.85 | Specific leaf area | 0.15 | 0.29 | 0.52 | 0.62 |
| | | | | | | Leaf water content | 0.21 | 0.36 | 0.57 | 0.59 |
| Gas exchange | 0.62 | 8.21 | 2 | 7 | **0.01** | Photosynthesis | 0.37 | 0.15 | 2.55 | **0.04** |
| | | | | | | Conductance | -0.34 | 0.08 | -4.03 | **0.005** |
| Phenology | 0.20 | 3.30 | 1 | 8 | 0.11 | Flowering time | -0.41 | 0.22 | -1.82 | 0.11 |

Bold p-values represent P<0.05. dfn = Degrees of freedom of numerator. dfd = Degrees of freed of denominator. Coeff = coefficient in the linear model. Predictor variables are evolution in each trait, estimated from slopes of trait values over time in a greenhouse common garden resurrection experiment (Anstett et al. 2021).

**Table S8. Ordinary least squares and robust regression for response to selection and nucleotide diversity predicting mean intrinsic rate of population growth during the recovery period (2016-18).**

| Comparison | Regression | Slope | SE | Adj $R^2$ | P-value | Figure |
|---|---|---|---|---|---|---|
| Response to Selection vs. Demographic Recovery | OLS | 6.60 | 2.01 | 0.52 | **0.011** | Fig. 3C |
| | Robust | 7.02 | 2.24 | NA | **0.012** | Fig. 3C |
| Climate-Associated PI vs. Demographic Recovery | OLS | 33.25 | 10.04 | 0.38 | **0.005** | Fig. 3D |
| | Robust | 35.13 | 10.23 | NA | **0.004** | Fig. 3D |
| Genome-Wide PI vs. Demographic Recovery | OLS | -8.34 | 13.78 | -0.04 | 0.5542 | Fig. 3E |
| | Robust | 1.02 | 6.18 | NA | 0.861 | Fig. 3E |

SE = Standard Error. Bold p-values represent P<0.05.

**Table S9. Vital rate models used to build integral projection models of population dynamics**.

| Vital rate | N | Model | Random effects structure | Family | R function (package) |
|---|---|---|---|---|---|
| Prob(Survival) | 19963 | logit(s) = a + b * z | (1\|Year/Site) | Binomial | glmer (lme4) |
| Growth | 5169 | z' = a + b * z | (logSize\|Year/Site) | Guassian | lmer (lme4) |
| Prob(Flowering) | 19963 | logit(fl) = a + b * z | (1\|Year/Site) | Binomial | glmer (lme4) |
| Fruit number | 3979 | log(fr) = a + b * z | (logSize\|Year) + (logSize\|Site) | Negative binomial | glmmTMB (glmmTMB) |

a = intercept, b = coefficient describing slope of vital rate with z, log(total stem length).

**Table S10. Vital rate coefficients (intercepts, slopes, and means and standard deviations (SD) when applicable) for integral projection modeling and resulting lambda estimates for each site and annual projection year. Prior to analyses, lambda values were converted to r = log(lambda + 0.01).**

| | | | | | | Prob (Survival) | | Growth | | | Prob (Flowering) | |
|---|---|---|---|---|---|---|---|---|---|---|---|---|
| **Population** | **Year** | **Latitude** | **Longitude** | **Elevation** | **Lambda** | **Intercept** | **Slope** | **Intercept** | **Slope** | **SD** | **Intercept** | **Slope** |
| Hauser Creek | 2010 | 32.65822 | -116.5323 | 788 | 0.1 | -4.46 | 0.5 | 1.94 | 0.49 | 1.1 | -6.15 | 1.85 |
| Hauser Creek | 2011 | 32.65822 | -116.5323 | 788 | 0.22 | -3.38 | 0.5 | 2.05 | 0.51 | 1.1 | -7.28 | 1.85 |
| Hauser Creek | 2012 | 32.65822 | -116.5323 | 788 | 0 | NA | NA | NA | NA | NA | NA | NA |
| Hauser Creek | 2013 | 32.65822 | -116.5323 | 788 | 0 | NA | NA | NA | NA | NA | NA | NA |
| Hauser Creek | 2014 | 32.65822 | -116.5323 | 788 | 0 | NA | NA | NA | NA | NA | NA | NA |
| Hauser Creek | 2015 | 32.65822 | -116.5323 | 788 | 0 | NA | NA | NA | NA | NA | NA | NA |
| Hauser Creek | 2016 | 32.65822 | -116.5323 | 788 | 0 | NA | NA | NA | NA | NA | NA | NA |
| Hauser Creek | 2017 | 32.65822 | -116.5323 | 788 | 0 | NA | NA | NA | NA | NA | NA | NA |
| Hauser Creek | 2018 | 32.65822 | -116.5323 | 788 | 1.95 | -3.75 | 0.5 | 2.17 | 0.53 | 1.1 | -5.5 | 1.85 |
| Kitchen Creek | 2010 | 32.75206 | -116.4522 | 1173 | 0.77 | -2.61 | 0.5 | 1.49 | 0.54 | 1.1 | -6.7 | 1.85 |
| Kitchen Creek | 2011 | 32.75206 | -116.4522 | 1173 | 0.85 | -3.23 | 0.5 | 1.41 | 0.61 | 1.1 | -7.58 | 1.85 |
| Kitchen Creek | 2012 | 32.75206 | -116.4522 | 1173 | 0.17 | -4.58 | 0.5 | 2.23 | 0.56 | 1.1 | -7.01 | 1.85 |
| Kitchen Creek | 2013 | 32.75206 | -116.4522 | 1173 | 0 | NA | NA | NA | NA | NA | NA | NA |
| Kitchen Creek | 2014 | 32.75206 | -116.4522 | 1173 | 0 | NA | NA | NA | NA | NA | NA | NA |
| Kitchen Creek | 2015 | 32.75206 | -116.4522 | 1173 | 0 | NA | NA | NA | NA | NA | NA | NA |
| Kitchen Creek | 2016 | 32.75206 | -116.4522 | 1173 | 0 | NA | NA | NA | NA | NA | NA | NA |
| Kitchen Creek | 2017 | 32.75206 | -116.4522 | 1173 | 0 | NA | NA | NA | NA | NA | NA | NA |
| Kitchen Creek | 2018 | 32.75206 | -116.4522 | 1173 | NA | NA | NA | NA | NA | NA | NA | NA |
| Sweetwater River | 2010 | 32.89913 | -116.5865 | 1386 | 0.61 | -2.92 | 0.5 | 2.06 | 0.66 | 1.1 | -7.21 | 1.85 |
| Sweetwater River | 2011 | 32.89913 | -116.5865 | 1386 | 0.51 | -2.49 | 0.5 | 1.71 | 0.6 | 1.1 | -7.71 | 1.85 |
| Sweetwater River | 2012 | 32.89913 | -116.5865 | 1386 | 0.29 | -3.22 | 0.5 | 1.99 | 0.5 | 1.1 | -7.82 | 1.85 |
| Sweetwater River | 2013 | 32.89913 | -116.5865 | 1386 | 0.64 | -2.1 | 0.5 | 2.1 | 0.55 | 1.1 | -6.66 | 1.85 |

| | | | | | | | | | | | | |
|---|---|---|---|---|---|---|---|---|---|---|---|---|
| Sweetwater River | 2014 | 32.89913 | -116.5865 | 1386 | 0.43 | -3.08 | 0.5 | 1.74 | 0.57 | 1.1 | -6.79 | 1.85 |
| Sweetwater River | 2015 | 32.89913 | -116.5865 | 1386 | 0.24 | -3.71 | 0.5 | 2.24 | 0.5 | 1.1 | -7.54 | 1.85 |
| Sweetwater River | 2016 | 32.89913 | -116.5865 | 1386 | 50.32 | -1.65 | 0.5 | 2.32 | 0.5 | 1.1 | -7.59 | 1.85 |
| Sweetwater River | 2017 | 32.89913 | -116.5865 | 1386 | 3.46 | -3.86 | 0.5 | 2.35 | 0.38 | 1.1 | -8.26 | 1.85 |
| Sweetwater River | 2018 | 32.89913 | -116.5865 | 1386 | 0.3 | -4.67 | 0.5 | 1.78 | 0.53 | 1.1 | -8.76 | 1.85 |
| Whitewater Canyon | 2010 | 33.99329 | -116.6627 | 696 | 0.04 | -5.75 | 0.5 | 1.79 | 0.54 | 1.1 | -3.32 | 1.85 |
| Whitewater Canyon | 2011 | 33.99329 | -116.6627 | 696 | 0.58 | -2.1 | 0.5 | 1.94 | 0.53 | 1.1 | -4.82 | 1.85 |
| Whitewater Canyon | 2012 | 33.99329 | -116.6627 | 696 | 0.54 | -2.53 | 0.5 | 1.64 | 0.59 | 1.1 | -6.19 | 1.85 |
| Whitewater Canyon | 2013 | 33.99329 | -116.6627 | 696 | 0.32 | -3.38 | 0.5 | 2.02 | 0.44 | 1.1 | -5.84 | 1.85 |
| Whitewater Canyon | 2014 | 33.99329 | -116.6627 | 696 | 0 | NA | NA | NA | NA | NA | NA | NA |
| Whitewater Canyon | 2015 | 33.99329 | -116.6627 | 696 | 0 | NA | NA | NA | NA | NA | NA | NA |
| Whitewater Canyon | 2016 | 33.99329 | -116.6627 | 696 | 0 | NA | NA | NA | NA | NA | NA | NA |
| Whitewater Canyon | 2017 | 33.99329 | -116.6627 | 696 | 0 | NA | NA | NA | NA | NA | NA | NA |
| Whitewater Canyon | 2018 | 33.99329 | -116.6627 | 696 | 0 | NA | NA | NA | NA | NA | NA | NA |
| West Fork Mojave River | 2010 | 34.28425 | -117.3754 | 1087 | 0.37 | -3.22 | 0.5 | 2.02 | 0.46 | 1.1 | -8.79 | 1.85 |
| West Fork Mojave River | 2011 | 34.28425 | -117.3754 | 1087 | 1.71 | -1.37 | 0.5 | 2.27 | 0.51 | 1.1 | -7.45 | 1.85 |
| West Fork Mojave River | 2012 | 34.28425 | -117.3754 | 1087 | 0.72 | -3.25 | 0.5 | 2.07 | 0.54 | 1.1 | -7.5 | 1.85 |
| West Fork Mojave River | 2013 | 34.28425 | -117.3754 | 1087 | NA | NA | NA | NA | NA | NA | NA | NA |
| West Fork Mojave River | 2014 | 34.28425 | -117.3754 | 1087 | 0.7 | -2.76 | 0.5 | 3.05 | 0.42 | 1.1 | -9.22 | 1.85 |
| West Fork Mojave River | 2015 | 34.28425 | -117.3754 | 1087 | 2.54 | -2.43 | 0.5 | 2.62 | 0.59 | 1.1 | -8.47 | 1.85 |
| West Fork Mojave River | 2016 | 34.28425 | -117.3754 | 1087 | 1.27 | -2.31 | 0.5 | 1.9 | 0.63 | 1.1 | -7 | 1.85 |
| West Fork Mojave River | 2017 | 34.28425 | -117.3754 | 1087 | 0.36 | -3.57 | 0.5 | 1.91 | 0.46 | 1.1 | -7.73 | 1.85 |
| West Fork Mojave River | 2018 | 34.28425 | -117.3754 | 1087 | 1.56 | -2.85 | 0.5 | 2.84 | 0.46 | 1.1 | -8.38 | 1.85 |
| South Fork Middle Fork Tule River | 2012 | 36.13756 | -118.5741 | 1664 | 1.64 | -1.42 | 0.5 | 2.34 | 0.59 | 1.1 | -8 | 1.85 |
| South Fork Middle Fork Tule River | 2013 | 36.13756 | -118.5741 | 1664 | 1.47 | -1.05 | 0.5 | 2.18 | 0.58 | 1.1 | -7.82 | 1.85 |
| South Fork Middle Fork Tule River | 2014 | 36.13756 | -118.5741 | 1664 | 0.87 | -1.41 | 0.5 | 2.51 | 0.43 | 1.1 | -7.82 | 1.85 |
| South Fork Middle Fork Tule River | 2015 | 36.13756 | -118.5741 | 1664 | 4.47 | -1.84 | 0.5 | 2.11 | 0.54 | 1.1 | -9.67 | 1.85 |

| | | | | | | | | | | | | |
|---|---|---|---|---|---|---|---|---|---|---|---|---|
| South Fork Middle Fork Tule River | 2018 | 36.13756 | -118.5741 | 1664 | 0.64 | -4.05 | 0.5 | 1.56 | 0.56 | 1.1 | -8.55 | 1.85 |
| North Fork Middle Fork Tule River | 2010 | 36.20081 | -118.6509 | 1284 | 0.89 | -2.97 | 0.5 | 2.36 | 0.51 | 1.1 | -10.24 | 1.85 |
| North Fork Middle Fork Tule | 2011 | 36.20081 | -118.6509 | 1284 | 0.81 | -1.89 | 0.5 | 2.27 | 0.53 | 1.1 | -9.21 | 1.85 |
| North Fork Middle Fork Tule River | 2012 | 36.20081 | -118.6509 | 1284 | 1.17 | -1.46 | 0.5 | 2.35 | 0.53 | 1.1 | -7.77 | 1.85 |
| North Fork Middle Fork Tule River | 2013 | 36.20081 | -118.6509 | 1284 | 1.03 | -1.39 | 0.5 | 0.89 | 0.82 | 1.1 | -7.59 | 1.85 |
| North Fork Middle Fork Tule River | 2014 | 36.20081 | -118.6509 | 1284 | 0.99 | -1.63 | 0.5 | 2.45 | 0.5 | 1.1 | -6.3 | 1.85 |
| North Fork Middle Fork Tule River | 2015 | 36.20081 | -118.6509 | 1284 | 1.42 | -2.47 | 0.5 | 2.16 | 0.53 | 1.1 | -7.86 | 1.85 |
| North Fork Middle Fork Tule River | 2018 | 36.20081 | -118.6509 | 1284 | 0.6 | -2.38 | 0.5 | 1.62 | 0.52 | 1.1 | -7.36 | 1.85 |
| Redwood Creek | 2010 | 36.68952 | -118.9096 | 1677 | 0.58 | -2.65 | 0.5 | 1.89 | 0.62 | 1.1 | -9.21 | 1.85 |
| Redwood Creek | 2011 | 36.68952 | -118.9096 | 1677 | 2.16 | -1.74 | 0.5 | 1.57 | 0.74 | 1.1 | -9.97 | 1.85 |
| Redwood Creek | 2012 | 36.68952 | -118.9096 | 1677 | 0.66 | -2.74 | 0.5 | 1.96 | 0.65 | 1.1 | -7.6 | 1.85 |
| Redwood Creek | 2013 | 36.68952 | -118.9096 | 1677 | 0.9 | -1.5 | 0.5 | 1.98 | 0.57 | 1.1 | -7.6 | 1.85 |
| Redwood Creek | 2014 | 36.68952 | -118.9096 | 1677 | 0.6 | -2.63 | 0.5 | 1.22 | 0.56 | 1.1 | -8.23 | 1.85 |
| Redwood Creek | 2016 | 36.68952 | -118.9096 | 1677 | NA | NA | NA | NA | NA | NA | NA | NA |
| Redwood Creek | 2017 | 36.68952 | -118.9096 | 1677 | 6.76 | -2.14 | 0.5 | 2.59 | 0.59 | 1.1 | -8.14 | 1.85 |
| Redwood Creek | 2018 | 36.68952 | -118.9096 | 1677 | 0.73 | -2.64 | 0.5 | 2.09 | 0.49 | 1.1 | -6.43 | 1.85 |
| Wawona | 2010 | 37.53822 | -119.6514 | 1208 | 1.19 | -1.08 | 0.5 | 2 | 0.6 | 1.1 | -7.91 | 1.85 |
| Wawona | 2011 | 37.53822 | -119.6514 | 1208 | 0.72 | -2.2 | 0.5 | 2.56 | 0.47 | 1.1 | -7.85 | 1.85 |
| Wawona | 2012 | 37.53822 | -119.6514 | 1208 | 0.91 | -2.32 | 0.5 | 2.48 | 0.51 | 1.1 | -7.5 | 1.85 |
| Wawona | 2013 | 37.53822 | -119.6514 | 1208 | 0.9 | -1.78 | 0.5 | 1.96 | 0.68 | 1.1 | -7.78 | 1.85 |
| Wawona | 2014 | 37.53822 | -119.6514 | 1208 | 0.8 | -2.25 | 0.5 | 2.93 | 0.5 | 1.1 | -7.44 | 1.85 |
| Wawona | 2015 | 37.53822 | -119.6514 | 1208 | 2.06 | -2.38 | 0.5 | 2.19 | 0.53 | 1.1 | -7.32 | 1.85 |
| Wawona | 2016 | 37.53822 | -119.6514 | 1208 | 1.05 | -2.19 | 0.5 | 2.54 | 0.48 | 1.1 | -7.83 | 1.85 |
| Wawona | 2017 | 37.53822 | -119.6514 | 1208 | 1.75 | -2.63 | 0.5 | 2.26 | 0.57 | 1.1 | -9.67 | 1.85 |
| Wawona | 2018 | 37.53822 | -119.6514 | 1208 | 0.74 | -2.44 | 0.5 | 1.98 | 0.57 | 1.1 | -8.08 | 1.85 |
| Buck Meadows | 2010 | 37.77722 | -120.0633 | 830 | 1.31 | -1.53 | 0.5 | 2.21 | 0.6 | 1.1 | -8.34 | 1.85 |

| | | | | | | | | | | | | |
|---|---|---|---|---|---|---|---|---|---|---|---|---|
| Buck Meadows | 2011 | 37.77722 | -120.0633 | 830 | 0.89 | -1.91 | 0.5 | 3.09 | 0.38 | 1.1 | -7.79 | 1.85 |
| Buck Meadows | 2014 | 37.77722 | -120.0633 | 830 | 0.53 | -2.24 | 0.5 | 2.58 | 0.42 | 1.1 | -8.12 | 1.85 |
| Buck Meadows | 2015 | 37.77722 | -120.0633 | 830 | 62.12 | -1.84 | 0.5 | 1.89 | 0.55 | 1.1 | -8.94 | 1.85 |
| Buck Meadows | 2016 | 37.77722 | -120.0633 | 830 | 5.6 | -3.42 | 0.5 | 2.55 | 0.59 | 1.1 | -8.09 | 1.85 |
| Buck Meadows | 2017 | 37.77722 | -120.0633 | 830 | 2.62 | -3.47 | 0.5 | 2.1 | 0.58 | 1.1 | -8.06 | 1.85 |
| Buck Meadows | 2018 | 37.77722 | -120.0633 | 830 | 0.72 | -2.13 | 0.5 | 1.65 | 0.55 | 1.1 | -9.62 | 1.85 |
| Carlon | 2010 | 37.81006 | -119.8568 | 1320 | 1 | -1.68 | 0.5 | 2.65 | 0.48 | 1.1 | -8.32 | 1.85 |
| Carlon | 2011 | 37.81006 | -119.8568 | 1320 | 0.3 | -3.55 | 0.5 | 2.44 | 0.25 | 1.1 | -8.87 | 1.85 |
| Carlon | 2014 | 37.81006 | -119.8568 | 1320 | 1.03 | -1.85 | 0.5 | 2.77 | 0.5 | 1.1 | -7.47 | 1.85 |
| Carlon | 2015 | 37.81006 | -119.8568 | 1320 | 0.48 | -3.24 | 0.5 | 2.24 | 0.55 | 1.1 | -8.53 | 1.85 |
| Carlon | 2016 | 37.81006 | -119.8568 | 1320 | 0.29 | -4.83 | 0.5 | 2.35 | 0.48 | 1.1 | -8.34 | 1.85 |
| Carlon | 2017 | 37.81006 | -119.8568 | 1320 | 10.8 | -3.57 | 0.5 | 1.97 | 0.55 | 1.1 | -8.25 | 1.85 |
| Carlon | 2018 | 37.81006 | -119.8568 | 1320 | 0.87 | -1.89 | 0.5 | 2.05 | 0.57 | 1.1 | -8.6 | 1.85 |
| Rainbow Pool | 2010 | 37.81878 | -120.0073 | 833 | 1.14 | -2.33 | 0.5 | 2.3 | 0.55 | 1.1 | -7.89 | 1.85 |
| Rainbow Pool | 2011 | 37.81878 | -120.0073 | 833 | 0.95 | -1.38 | 0.5 | 2.36 | 0.53 | 1.1 | -6.69 | 1.85 |
| Rainbow Pool | 2014 | 37.81878 | -120.0073 | 833 | 0.8 | -2.25 | 0.5 | 1.69 | 0.74 | 1.1 | -6.19 | 1.85 |
| Rainbow Pool | 2015 | 37.81878 | -120.0073 | 833 | 0.85 | -3.34 | 0.5 | 1.8 | 0.59 | 1.1 | -7.24 | 1.85 |
| Rainbow Pool | 2016 | 37.81878 | -120.0073 | 833 | 1.48 | -3.04 | 0.5 | 1.91 | 0.58 | 1.1 | -7.83 | 1.85 |
| Rainbow Pool | 2017 | 37.81878 | -120.0073 | 833 | 7.64 | -2.31 | 0.5 | 2.17 | 0.5 | 1.1 | -6.63 | 1.85 |
| Rainbow Pool | 2018 | 37.81878 | -120.0073 | 833 | 2.86 | -2.03 | 0.5 | 1.77 | 0.66 | 1.1 | -6.83 | 1.85 |
| Oregon Creek | 2010 | 39.39704 | -121.082 | 448 | 1.43 | -2.32 | 0.5 | 2.4 | 0.52 | 1.1 | -7.83 | 1.85 |
| Oregon Creek | 2011 | 39.39704 | -121.082 | 448 | 1.21 | -2.56 | 0.5 | 2.27 | 0.41 | 1.1 | -8.26 | 1.85 |
| Oregon Creek | 2012 | 39.39704 | -121.082 | 448 | 1.72 | -1.11 | 0.5 | 1.98 | 0.57 | 1.1 | -8.15 | 1.85 |
| Oregon Creek | 2013 | 39.39704 | -121.082 | 448 | 1.29 | -1.08 | 0.5 | 2.32 | 0.49 | 1.1 | -8.01 | 1.85 |
| Oregon Creek | 2014 | 39.39704 | -121.082 | 448 | 0.98 | -1.69 | 0.5 | 2.07 | 0.55 | 1.1 | -7.6 | 1.85 |
| Oregon Creek | 2015 | 39.39704 | -121.082 | 448 | 0.21 | -4.19 | 0.5 | 1.5 | 0.5 | 1.1 | -8.67 | 1.85 |
| Oregon Creek | 2016 | 39.39704 | -121.082 | 448 | 4.2 | -3.11 | 0.5 | 1.85 | 0.57 | 1.1 | -8.56 | 1.85 |
| Oregon Creek | 2017 | 39.39704 | -121.082 | 448 | 6.28 | -1.76 | 0.5 | 2.34 | 0.58 | 1.1 | -7.43 | 1.85 |
| Oregon Creek | 2018 | 39.39704 | -121.082 | 448 | 1.57 | -1.16 | 0.5 | 2.19 | 0.56 | 1.1 | -7.22 | 1.85 |

| | | | | | | | | | | | | |
|---|---|---|---|---|---|---|---|---|---|---|---|---|
| Little Jamison Creek | 2010 | 39.74298 | -120.704 | 1592 | 1.3 | -0.87 | 0.5 | 2.66 | 0.65 | 1.1 | -8.05 | 1.85 |
| Little Jamison Creek | 2011 | 39.74298 | -120.704 | 1592 | 1.04 | -1.54 | 0.5 | 1.93 | 0.82 | 1.1 | -8.92 | 1.85 |
| Little Jamison Creek | 2012 | 39.74298 | -120.704 | 1592 | 0.76 | -2.05 | 0.5 | 1.98 | 0.67 | 1.1 | -7.93 | 1.85 |
| Little Jamison Creek | 2013 | 39.74298 | -120.704 | 1592 | 0.83 | -1.81 | 0.5 | 2.19 | 0.66 | 1.1 | -6.61 | 1.85 |
| Little Jamison Creek | 2014 | 39.74298 | -120.704 | 1592 | 0.91 | -0.93 | 0.5 | 2.7 | 0.6 | 1.1 | -6.54 | 1.85 |
| Little Jamison Creek | 2015 | 39.74298 | -120.704 | 1592 | 0.78 | -2.04 | 0.5 | 1.82 | 0.68 | 1.1 | -6.87 | 1.85 |
| Little Jamison Creek | 2016 | 39.74298 | -120.704 | 1592 | 0.69 | -2.84 | 0.5 | 2.21 | 0.65 | 1.1 | -7.55 | 1.85 |
| Little Jamison Creek | 2017 | 39.74298 | -120.704 | 1592 | 1.11 | -1.56 | 0.5 | 2.56 | 0.63 | 1.1 | -8.26 | 1.85 |
| Little Jamison Creek | 2018 | 39.74298 | -120.704 | 1592 | 0.8 | -2.04 | 0.5 | 2.25 | 0.62 | 1.1 | -6.37 | 1.85 |
| Deep Creek | 2010 | 41.66546 | -123.1134 | 694 | 1.37 | -0.27 | 0.5 | 2.06 | 0.66 | 1.1 | -8.75 | 1.85 |
| Deep Creek | 2011 | 41.66546 | -123.1134 | 694 | 2.67 | -1.06 | 0.5 | 1.73 | 0.66 | 1.1 | -10.52 | 1.85 |
| Deep Creek | 2012 | 41.66546 | -123.1134 | 694 | 1.08 | -0.86 | 0.5 | 1.38 | 0.65 | 1.1 | -8.83 | 1.85 |
| Deep Creek | 2013 | 41.66546 | -123.1134 | 694 | 0.95 | -1.11 | 0.5 | 1.35 | 0.67 | 1.1 | -8.75 | 1.85 |
| Deep Creek | 2014 | 41.66546 | -123.1134 | 694 | 2.63 | -1.58 | 0.5 | 1.24 | 0.7 | 1.1 | -8.32 | 1.85 |
| Deep Creek | 2015 | 41.66546 | -123.1134 | 694 | 0.91 | -1.89 | 0.5 | 2.01 | 0.72 | 1.1 | -7.17 | 1.85 |
| Deep Creek | 2016 | 41.66546 | -123.1134 | 694 | 0.47 | -2.59 | 0.5 | 1.1 | 0.65 | 1.1 | -7.63 | 1.85 |
| Deep Creek | 2017 | 41.66546 | -123.1134 | 694 | 0.75 | -3 | 0.5 | 1.64 | 0.58 | 1.1 | -8.17 | 1.85 |
| Deep Creek | 2018 | 41.66546 | -123.1134 | 694 | 1.58 | -2.02 | 0.5 | 3.73 | 0.35 | 1.1 | -8.6 | 1.85 |
| O'Neil Creek | 2010 | 41.80979 | -123.1189 | 487 | 1.31 | -0.61 | 0.5 | 2.41 | 0.61 | 1.1 | -9.67 | 1.85 |
| O'Neil Creek | 2011 | 41.80979 | -123.1189 | 487 | 1.39 | -1.53 | 0.5 | 2.18 | 0.6 | 1.1 | -9.26 | 1.85 |
| O'Neil Creek | 2012 | 41.80979 | -123.1189 | 487 | 0.27 | -2.96 | 0.5 | 2.27 | 0.31 | 1.1 | -9.96 | 1.85 |
| O'Neil Creek | 2013 | 41.80979 | -123.1189 | 487 | 3.34 | -1.82 | 0.5 | 2.76 | 0.47 | 1.1 | -10.26 | 1.85 |
| O'Neil Creek | 2014 | 41.80979 | -123.1189 | 487 | 2.18 | -2.18 | 0.5 | 2.57 | 0.42 | 1.1 | -9.59 | 1.85 |
| O'Neil Creek | 2015 | 41.80979 | -123.1189 | 487 | 3.55 | -0.76 | 0.5 | 2.99 | 0.55 | 1.1 | -6.37 | 1.85 |
| O'Neil Creek | 2016 | 41.80979 | -123.1189 | 487 | 0.27 | -4.15 | 0.5 | 2.62 | 0.35 | 1.1 | -7.28 | 1.85 |
| O'Neil Creek | 2017 | 41.80979 | -123.1189 | 487 | 3.09 | -1.4 | 0.5 | 3.52 | 0.52 | 1.1 | -8.05 | 1.85 |
| O'Neil Creek | 2018 | 41.80979 | -123.1189 | 487 | 0.72 | -2.09 | 0.5 | 2.29 | 0.54 | 1.1 | -7.31 | 1.85 |
| Rock Creek | 2010 | 43.37876 | -122.9521 | 295 | 0.17 | -4.48 | 0.5 | 1.87 | 0.53 | 1.1 | -8.82 | 1.85 |
| Rock Creek | 2011 | 43.37876 | -122.9521 | 295 | 0.73 | -3.42 | 0.5 | 3.2 | 0.51 | 1.1 | -8.71 | 1.85 |

| | | | | | | | | | | | | |
|---|---|---|---|---|---|---|---|---|---|---|---|---|
| Rock Creek | 2014 | 43.37876 | -122.9521 | 295 | 0.34 | -4.58 | 0.5 | 1.11 | 0.57 | 1.1 | -8.1 | 1.85 |
| Rock Creek | 2015 | 43.37876 | -122.9521 | 295 | 0.52 | -3.52 | 0.5 | 2.56 | 0.53 | 1.1 | -5.71 | 1.85 |
| Rock Creek | 2016 | 43.37876 | -122.9521 | 295 | 4.26 | -2.29 | 0.5 | 2.76 | 0.46 | 1.1 | -6.2 | 1.85 |
| Rock Creek | 2017 | 43.37876 | -122.9521 | 295 | 5.61 | -0.48 | 0.5 | 2.25 | 0.57 | 1.1 | -7.23 | 1.85 |
| Rock Creek | 2018 | 43.37876 | -122.9521 | 295 | 0.69 | -2.81 | 0.5 | 2.07 | 0.44 | 1.1 | -7.24 | 1.85 |
| Canton Creek | 2010 | 43.4137 | -122.7804 | 503 | 2.86 | -1.91 | 0.5 | 2.42 | 0.49 | 1.1 | -10.18 | 1.85 |
| Canton Creek | 2011 | 43.4137 | -122.7804 | 503 | 10.46 | -4.92 | 0.5 | 2.4 | 0.49 | 1.1 | -10.12 | 1.85 |
| Canton Creek | 2012 | 43.4137 | -122.7804 | 503 | 3.56 | -1.57 | 0.5 | 3.21 | 0.32 | 1.1 | -8.45 | 1.85 |
| Canton Creek | 2013 | 43.4137 | -122.7804 | 503 | 0.89 | -1.91 | 0.5 | 2.79 | 0.54 | 1.1 | -7.84 | 1.85 |
| Canton Creek | 2014 | 43.4137 | -122.7804 | 503 | 0.5 | -3.34 | 0.5 | 1.8 | 0.59 | 1.1 | -8.42 | 1.85 |
| Canton Creek | 2015 | 43.4137 | -122.7804 | 503 | 1.09 | -1.41 | 0.5 | 1.83 | 0.57 | 1.1 | -8.62 | 1.85 |
| Canton Creek | 2017 | 43.4137 | -122.7804 | 503 | NA | NA | NA | NA | NA | NA | NA | NA |
| Canton Creek | 2018 | 43.4137 | -122.7804 | 503 | 0.5 | -2.33 | 0.5 | 2.38 | 0.43 | 1.1 | -8.77 | 1.85 |
| Coast Fork Willamette River | 2010 | 43.6517 | -123.088 | 254 | 1.73 | -0.6 | 0.5 | 2.18 | 0.54 | 1.1 | -7.71 | 1.85 |
| Coast Fork Willamette River | 2011 | 43.6517 | -123.088 | 254 | 7.04 | -3.74 | 0.5 | 2.18 | 0.55 | 1.1 | -8.71 | 1.85 |
| Coast Fork Willamette River | 2012 | 43.6517 | -123.088 | 254 | 12.32 | -2.08 | 0.5 | 2.6 | 0.54 | 1.1 | -8.83 | 1.85 |
| Coast Fork Willamette River | 2013 | 43.6517 | -123.088 | 254 | 0.62 | -3.04 | 0.5 | 2.67 | 0.57 | 1.1 | -7.61 | 1.85 |
| Coast Fork Willamette River | 2014 | 43.6517 | -123.088 | 254 | 1.05 | -2.54 | 0.5 | 2.29 | 0.6 | 1.1 | -7.97 | 1.85 |
| Coast Fork Willamette River | 2015 | 43.6517 | -123.088 | 254 | 0.66 | -2.7 | 0.5 | 3.23 | 0.38 | 1.1 | -7.03 | 1.85 |
| Coast Fork Willamette River | 2016 | 43.6517 | -123.088 | 254 | 0.58 | -2.92 | 0.5 | 2.13 | 0.5 | 1.1 | -8.17 | 1.85 |
| Coast Fork Willamette River | 2017 | 43.6517 | -123.088 | 254 | 7.92 | -2 | 0.5 | 2.68 | 0.48 | 1.1 | -7.83 | 1.85 |
| Coast Fork Willamette River | 2018 | 43.6517 | -123.088 | 254 | 1.95 | -3.51 | 0.5 | 2.32 | 0.38 | 1.1 | -7.1 | 1.85 |

| | | **Fruit Number** | | **Seeds per Fruit** | **Prob (Recruitment)** | **Distribution of Recruit Size** | |
|---|---|---|---|---|---|---|---|
| **Population** | **Year** | **Intercept** | **Slope** | | | **Mean** | **SD** |
| Hauser Creek | 2010 | 0.4 | 0.57 | 1666 | 0 | 0 | 0 |
| Hauser Creek | 2011 | 0.94 | 0.39 | 1666 | 0 | 0 | 0 |

| | | | | | | | |
|---|---|---|---|---|---|---|---|
| Hauser Creek | 2012 | NA | NA | NA | NA | NA | NA |
| Hauser Creek | 2013 | NA | NA | NA | NA | NA | NA |
| Hauser Creek | 2014 | NA | NA | NA | NA | NA | NA |
| Hauser Creek | 2015 | NA | NA | NA | NA | NA | NA |
| Hauser Creek | 2016 | NA | NA | NA | NA | NA | NA |
| Hauser Creek | 2017 | NA | NA | NA | NA | NA | NA |
| Hauser Creek | 2018 | 0.01 | 0.66 | 1666 | 0.000436861 | 2.33 | 0.73 |
| Kitchen Creek | 2010 | -0.78 | 0.83 | 2514 | 0.000169265 | 1.17 | 1.25 |
| Kitchen Creek | 2011 | -0.24 | 0.65 | 2514 | 0.000318218 | 1.73 | 1.36 |
| Kitchen Creek | 2012 | -1.11 | 0.88 | 2514 | 0 | 0 | 0 |
| Kitchen Creek | 2013 | NA | NA | NA | NA | NA | NA |
| Kitchen Creek | 2014 | NA | NA | NA | NA | NA | NA |
| Kitchen Creek | 2015 | NA | NA | NA | NA | NA | NA |
| Kitchen Creek | 2016 | NA | NA | NA | NA | NA | NA |
| Kitchen Creek | 2017 | NA | NA | NA | NA | NA | NA |
| Kitchen Creek | 2018 | NA | NA | NA | NA | NA | NA |
| Sweetwater River | 2010 | 0.07 | 0.59 | 2027 | 9.02E-07 | 3.64 | 0.8 |
| Sweetwater River | 2011 | 0.61 | 0.42 | 2027 | 1.38E-06 | 2.77 | 1.4 |
| Sweetwater River | 2012 | -0.26 | 0.64 | 2027 | 2.78E-07 | 2.21 | 0.37 |
| Sweetwater River | 2013 | -0.02 | 0.61 | 2027 | 2.51E-06 | 2.77 | 1.1 |
| Sweetwater River | 2014 | -0.13 | 0.61 | 2027 | 2.26E-05 | 2.37 | 0.6 |
| Sweetwater River | 2015 | 0.17 | 0.53 | 2027 | 8.22E-05 | 2.64 | 0 |
| Sweetwater River | 2016 | -0.54 | 0.74 | 2027 | 0.086968779 | 1.35 | 1.14 |
| Sweetwater River | 2017 | -0.3 | 0.64 | 2027 | 0.000523544 | 3.13 | 1.24 |
| Sweetwater River | 2018 | -0.32 | 0.69 | 2027 | 0.000733484 | 1.26 | 0.89 |
| Whitewater Canyon | 2010 | 1.37 | 0.43 | 1104 | 3.95E-07 | 0 | 0 |
| Whitewater Canyon | 2011 | 1.91 | 0.26 | 1104 | 5.83E-06 | 2.39 | 0.97 |
| Whitewater Canyon | 2012 | 1.04 | 0.49 | 1104 | 4.09E-06 | 4.08 | 1.05 |
| Whitewater Canyon | 2013 | 1.29 | 0.46 | 1104 | 4.66E-06 | 3.98 | 1.02 |

| | | | | | | | |
|---|---|---|---|---|---|---|---|
| Whitewater Canyon | 2014 | NA | NA | NA | NA | NA | NA |
| Whitewater Canyon | 2015 | NA | NA | NA | NA | NA | NA |
| Whitewater Canyon | 2016 | NA | NA | NA | NA | NA | NA |
| Whitewater Canyon | 2017 | NA | NA | NA | NA | NA | NA |
| Whitewater Canyon | 2018 | NA | NA | NA | NA | NA | NA |
| West Fork Mojave River | 2010 | 0.78 | 0.42 | 1675 | 1.84E-05 | 3.77 | 0.39 |
| West Fork Mojave River | 2011 | 1.32 | 0.24 | 1675 | 0.000118492 | 3.65 | 0.88 |
| West Fork Mojave River | 2012 | 0.45 | 0.47 | 1675 | 2.84E-05 | 4.15 | 0.93 |
| West Fork Mojave River | 2013 | NA | NA | NA | NA | NA | NA |
| West Fork Mojave River | 2014 | 0.59 | 0.44 | 1675 | 3.57E-05 | 3.17 | 0.62 |
| West Fork Mojave River | 2015 | 0.88 | 0.36 | 1675 | 0.000526778 | 1.84 | 2.05 |
| West Fork Mojave River | 2016 | 0.17 | 0.56 | 1675 | 6.35E-06 | 7.22 | 1.4 |
| West Fork Mojave River | 2017 | 0.41 | 0.46 | 1675 | 5.54E-06 | 5 | 1.89 |
| West Fork Mojave River | 2018 | 0.39 | 0.51 | 1675 | 0.000523333 | 1.27 | 1.68 |
| South Fork Middle Fork Tule River | 2012 | 0.07 | 0.51 | 1473 | 0.000382071 | 1.54 | 1.08 |
| South Fork Middle Fork Tule River | 2013 | 0.31 | 0.48 | 1473 | 0.000452591 | 0.41 | 1.28 |
| South Fork Middle Fork Tule River | 2014 | 0.2 | 0.48 | 1473 | 4.61E-05 | 1.53 | 1.44 |
| South Fork Middle Fork Tule River | 2015 | 0.49 | 0.4 | 1473 | 0.001804845 | 2.78 | 1.59 |
| South Fork Middle Fork Tule River | 2018 | 0.01 | 0.56 | 1473 | 0.000274345 | 2.11 | 1.52 |
| North Fork Middle Fork Tule River | 2010 | 0.54 | 0.47 | 1473 | 0.003281285 | 0.14 | 1.1 |
| North Fork Middle Fork Tule River | 2011 | 1.07 | 0.29 | 1473 | 0.000307115 | -0.04 | 0.83 |
| North Fork Middle Fork Tule River | 2012 | 0.21 | 0.52 | 1473 | 0.000228471 | 0.62 | 1.17 |
| North Fork Middle Fork Tule River | 2013 | 0.45 | 0.49 | 1473 | 0.000174865 | 0.92 | 1.04 |
| North Fork Middle Fork Tule River | 2014 | 0.34 | 0.49 | 1473 | 7.77E-05 | 0.97 | 1.43 |

| | | | | | | | |
|---|---|---|---|---|---|---|---|
| North Fork Middle Fork Tule River | 2015 | 0.64 | 0.41 | 1473 | 0.002095693 | 0.67 | 1.24 |
| North Fork Middle Fork Tule River | 2018 | 0.15 | 0.57 | 1473 | 0.000194423 | 0.78 | 1.23 |
| Redwood Creek | 2010 | 0.71 | 0.44 | 1289 | 0.000113245 | -0.04 | 1.18 |
| Redwood Creek | 2011 | 1.25 | 0.27 | 1289 | 0.001388265 | 2.35 | 1.28 |
| Redwood Creek | 2012 | 0.38 | 0.49 | 1289 | 3.20E-05 | 1.35 | 1.36 |
| Redwood Creek | 2013 | 0.63 | 0.46 | 1289 | 0.000183426 | -0.11 | 0.92 |
| Redwood Creek | 2014 | 0.52 | 0.46 | 1289 | 0.000286898 | 1.93 | 1.16 |
| Redwood Creek | 2016 | NA | NA | NA | NA | NA | NA |
| Redwood Creek | 2017 | 0.35 | 0.49 | 1289 | 0.001393265 | 3.27 | 1.12 |
| Redwood Creek | 2018 | 0.33 | 0.54 | 1289 | 5.84E-05 | 2.08 | 1.7 |
| Wawona | 2010 | -0.21 | 0.62 | 1721 | 0.000280951 | -0.23 | 1 |
| Wawona | 2011 | 0.33 | 0.44 | 1721 | 9.34E-05 | 0.39 | 1.2 |
| Wawona | 2012 | -0.54 | 0.67 | 1721 | 0.000256208 | 0.37 | 1.09 |
| Wawona | 2013 | -0.29 | 0.64 | 1721 | 3.34E-05 | 0.17 | 1.22 |
| Wawona | 2014 | -0.4 | 0.64 | 1721 | 7.73E-06 | 2.44 | 1.54 |
| Wawona | 2015 | -0.11 | 0.56 | 1721 | 0.000222989 | 2.16 | 2.11 |
| Wawona | 2016 | -0.82 | 0.76 | 1721 | 0.000178534 | 0.93 | 1.37 |
| Wawona | 2017 | -0.58 | 0.66 | 1721 | 0.00040674 | 2.94 | 1.1 |
| Wawona | 2018 | -0.6 | 0.71 | 1721 | 2.29E-05 | 2.35 | 1.67 |
| Buck Meadows | 2010 | 1.62 | 0.13 | 1956 | 0.000529506 | 0.75 | 1.25 |
| Buck Meadows | 2011 | 2.16 | -0.05 | 1956 | 7.53E-05 | 2.41 | 1.2 |
| Buck Meadows | 2014 | 1.43 | 0.15 | 1956 | 1.55E-05 | 4.51 | 0 |
| Buck Meadows | 2015 | 1.72 | 0.07 | 1956 | 0.107617587 | 2.04 | 1.25 |
| Buck Meadows | 2016 | 1.02 | 0.27 | 1956 | 0.008918428 | 1.56 | 1.21 |
| Buck Meadows | 2017 | 1.26 | 0.18 | 1956 | 0.000471484 | 3.54 | 1.28 |
| Buck Meadows | 2018 | 1.24 | 0.22 | 1956 | 0.000261825 | 1.78 | 1.24 |
| Carlon | 2010 | -0.68 | 0.7 | 1226 | 4.14E-05 | 2.05 | 1.65 |
| Carlon | 2011 | -0.14 | 0.52 | 1226 | 0.000139903 | 1.93 | 1.62 |

| | | | | | | | |
|---|---|---|---|---|---|---|---|
| Carlon | 2014 | -0.87 | 0.72 | 1226 | 3.40E-05 | 2.59 | 1.22 |
| Carlon | 2015 | -0.58 | 0.64 | 1226 | 2.58E-05 | 2.36 | 1.36 |
| Carlon | 2016 | -1.29 | 0.84 | 1226 | 0.000286011 | 1.97 | 0.89 |
| Carlon | 2017 | -1.05 | 0.75 | 1226 | 0.01223491 | 2.39 | 1.14 |
| Carlon | 2018 | -1.07 | 0.8 | 1226 | 7.42E-05 | 1.63 | 0.66 |
| Rainbow Pool | 2010 | 1.21 | 0.32 | 2144 | 0.000156356 | 1.58 | 1.56 |
| Rainbow Pool | 2011 | 1.75 | 0.14 | 2144 | 7.54E-05 | 0.23 | 1.39 |
| Rainbow Pool | 2014 | 1.01 | 0.34 | 2144 | 3.99E-05 | 0.63 | 1.34 |
| Rainbow Pool | 2015 | 1.31 | 0.26 | 2144 | 0.000514161 | 1.38 | 1.08 |
| Rainbow Pool | 2016 | 0.6 | 0.46 | 2144 | 0.001700169 | 1.18 | 1.07 |
| Rainbow Pool | 2017 | 0.84 | 0.37 | 2144 | 0.003524046 | 1.94 | 0.98 |
| Rainbow Pool | 2018 | 0.82 | 0.42 | 2144 | 0.000660759 | 2.02 | 1.07 |
| Oregon Creek | 2010 | 0.28 | 0.57 | 1691 | 0.00019557 | 1.71 | 1.58 |
| Oregon Creek | 2011 | 0.82 | 0.4 | 1691 | 0.001960845 | 0.48 | 1.35 |
| Oregon Creek | 2012 | -0.05 | 0.63 | 1691 | 0.000381662 | 1.37 | 1.21 |
| Oregon Creek | 2013 | 0.2 | 0.6 | 1691 | 0.00019285 | 0.45 | 1.29 |
| Oregon Creek | 2014 | 0.09 | 0.59 | 1691 | 2.90E-05 | 2.85 | 1 |
| Oregon Creek | 2015 | 0.38 | 0.51 | 1691 | 0.000128558 | 1.75 | 1.13 |
| Oregon Creek | 2016 | -0.33 | 0.72 | 1691 | 0.001669739 | 2.51 | 1.15 |
| Oregon Creek | 2017 | -0.08 | 0.62 | 1691 | 0.001492495 | 2.31 | 1.27 |
| Oregon Creek | 2018 | -0.11 | 0.67 | 1691 | 7.49E-05 | 2.34 | 1.23 |
| Little Jamison Creek | 2010 | -1.13 | 0.92 | 2092 | 6.10E-06 | 0.08 | 1.03 |
| Little Jamison Creek | 2011 | -0.59 | 0.74 | 2092 | 4.72E-07 | 1.54 | 1.29 |
| Little Jamison Creek | 2012 | -1.46 | 0.97 | 2092 | 7.90E-08 | 2.88 | 1.26 |
| Little Jamison Creek | 2013 | -1.21 | 0.94 | 2092 | 4.51E-08 | 3.39 | 2.06 |
| Little Jamison Creek | 2014 | -1.32 | 0.94 | 2092 | 0 | 0 | 0 |
| Little Jamison Creek | 2015 | -1.03 | 0.86 | 2092 | 4.15E-06 | 0.93 | 0.79 |
| Little Jamison Creek | 2016 | -1.74 | 1.06 | 2092 | 2.53E-06 | 0.86 | 1.3 |
| Little Jamison Creek | 2017 | -1.5 | 0.96 | 2092 | 7.35E-07 | 4.26 | 1.63 |

| | | | | | | | |
|---|---|---|---|---|---|---|---|
| Little Jamison Creek | 2018 | -1.52 | 1.01 | 2092 | 8.02E-07 | 1.5 | 1.66 |
| Deep Creek | 2010 | 0.91 | 0.41 | 506 | 5.25E-05 | 4.46 | 0.96 |
| Deep Creek | 2011 | 1.45 | 0.23 | 506 | 0.001288881 | 2.88 | 2.56 |
| Deep Creek | 2012 | 0.58 | 0.46 | 506 | 0.000228033 | 2.16 | 1.56 |
| Deep Creek | 2013 | 0.82 | 0.43 | 506 | 0.000658762 | -0.04 | 1.3 |
| Deep Creek | 2014 | 0.72 | 0.43 | 506 | 0.011749418 | 0.86 | 1.43 |
| Deep Creek | 2015 | 1.01 | 0.35 | 506 | 3.03E-05 | 1.88 | 1.81 |
| Deep Creek | 2016 | 0.3 | 0.55 | 506 | 3.52E-05 | 2.35 | 1.74 |
| Deep Creek | 2017 | 0.54 | 0.46 | 506 | 7.47E-05 | 4.12 | 1.78 |
| Deep Creek | 2018 | 0.52 | 0.51 | 506 | 0.00011074 | 4.09 | 1.9 |
| O'Neil Creek | 2010 | 0.39 | 0.53 | 1017 | 3.53E-05 | 2.2 | 2.32 |
| O'Neil Creek | 2011 | 0.93 | 0.35 | 1017 | 0.000146164 | 3.08 | 1.5 |
| O'Neil Creek | 2012 | 0.06 | 0.58 | 1017 | 2.25E-05 | 1.76 | 1.42 |
| O'Neil Creek | 2013 | 0.31 | 0.55 | 1017 | 0.001638807 | 3.29 | 0.77 |
| O'Neil Creek | 2014 | 0.2 | 0.55 | 1017 | 0.003241196 | 1.57 | 1.31 |
| O'Neil Creek | 2015 | 0.49 | 0.47 | 1017 | 0.001161504 | 0.95 | 1.69 |
| O'Neil Creek | 2016 | -0.22 | 0.67 | 1017 | 1.90E-05 | 3.32 | 1.18 |
| O'Neil Creek | 2017 | 0.02 | 0.58 | 1017 | 0.000144215 | 3.77 | 1.84 |
| O'Neil Creek | 2018 | 0 | 0.62 | 1017 | 6.64E-06 | 3.09 | 1.64 |
| Rock Creek | 2010 | 0.93 | 0.5 | 1122 | 1.41E-05 | 2.53 | 1.22 |
| Rock Creek | 2011 | 1.46 | 0.32 | 1122 | 0.000124263 | 1.97 | 1.07 |
| Rock Creek | 2014 | 0.73 | 0.52 | 1122 | 8.76E-05 | 2.71 | 1.17 |
| Rock Creek | 2015 | 1.03 | 0.44 | 1122 | 4.77E-05 | 1.88 | 1.03 |
| Rock Creek | 2016 | 0.32 | 0.64 | 1122 | 0.000926217 | 2.52 | 0.79 |
| Rock Creek | 2017 | 0.56 | 0.55 | 1122 | 0.001147842 | 2.52 | 1.08 |
| Rock Creek | 2018 | 0.54 | 0.6 | 1122 | 6.91E-05 | 2.83 | 0.77 |
| Canton Creek | 2010 | 0.65 | 0.41 | 1184 | 0.002252252 | 2.32 | 1.48 |
| Canton Creek | 2011 | 1.19 | 0.24 | 1184 | 0.027027027 | 2.16 | 1.48 |
| Canton Creek | 2012 | 0.32 | 0.47 | 1184 | 0.002051158 | 2.43 | 1.03 |

| | | | | | | | |
|---|---|---|---|---|---|---|---|
| Canton Creek | 2013 | 0.57 | 0.44 | 1184 | 2.93E-05 | 2.11 | 0.62 |
| Canton Creek | 2014 | 0.46 | 0.44 | 1184 | 0.000120656 | 2 | 1.48 |
| Canton Creek | 2015 | 0.75 | 0.36 | 1184 | 0.000469219 | 1.31 | 1.02 |
| Canton Creek | 2017 | NA | NA | NA | NA | NA | NA |
| Canton Creek | 2018 | 0.26 | 0.51 | 1184 | 3.82E-06 | 2.46 | 0.52 |
| Coast Fork Willamette | 2010 | 0.66 | 0.44 | 1184 | 0.000440658 | 1.48 | 1.01 |
| Coast Fork Willamette | 2011 | 1.19 | 0.26 | 1184 | 0.016047297 | 1.93 | 1.2 |
| Coast Fork Willamette | 2012 | 0.33 | 0.49 | 1184 | 0.077702703 | 1.37 | 1.04 |
| Coast Fork Willamette | 2013 | 0.57 | 0.46 | 1184 | 7.72E-05 | 1.33 | 0.83 |
| Coast Fork Willamette | 2014 | 0.46 | 0.46 | 1184 | 0.000224229 | 2.18 | 0.94 |
| Coast Fork Willamette | 2015 | 0.76 | 0.38 | 1184 | 3.54E-05 | 2.13 | 1.66 |
| Coast Fork Willamette | 2016 | 0.05 | 0.58 | 1184 | 4.33E-05 | 3.31 | 0.76 |
| Coast Fork Willamette | 2017 | 0.29 | 0.49 | 1184 | 0.004898649 | 2.38 | 1.17 |
| Coast Fork Willamette | 2018 | 0.27 | 0.54 | 1184 | 0.000499718 | 2.63 | 1.28 |

**Table S11. Statistics for median response to selection (S) on alleles associated with three climate variables across the 11 timeseries populations.**

| Population | Population ID | Latitude | Longitude | Median S | Wilcoxon P-value | Median Percentile | Median P-value |
|---|---|---|---|---|---|---|---|
| Sweetwater River | 1 | 32.89913 | −116.5865 | 0.05 | 0.07 | 0.83 | 0.17 |
| West Fork Mojave River | 2 | 34.28425 | −117.3754 | -0.06 | 0.43 | *0.074* | 0.93 |
| North Fork Middle Fork Tule River | 3 | 36.20081 | −118.6509 | 0.02 | 0.33 | 0.69 | 0.31 |
| Redwood Creek | 4 | 36.68952 | −118.9096 | 0.10 | 0.18 | 0.78 | 0.22 |
| Wawona | 5 | 37.53822 | −119.6514 | 0.00 | 0.51 | 0.55 | 0.45 |
| Oregon Creek | 6 | 39.39704 | −121.082 | 0.00 | 0.41 | 0.50 | 0.50 |
| Little Jamison Creek | 7 | 39.74298 | −120.704 | -0.09 | 0.00 | **0.005** | 1.00 |
| Deep Creek | 8 | 41.66546 | −123.1134 | -0.07 | 0.17 | **0.035** | 0.97 |

| | | | | | | | |
|---|---|---|---|---|---|---|---|
| O'Neil Creek | 9 | 41.80979 | −123.1189 | 0.03 | 0.26 | 0.76 | 0.24 |
| Deer Creek | 10 | 42.27411 | −123.63617 | -0.02 | 0.26 | 0.25 | 0.75 |
| Rock Creek | 11 | 43.37876 | −122.9521 | 0.08 | 0.02 | 0.99 | **0.01** |

Median S is the median climate-associated response to selection, derived from logistic regression of allele frequency across years. Wilcoxon p-value is given for the test that the median S of climate-associated loci is different from 0. Sets of non-climate associated SNPs were randomly selected 1000 times to generate 1000 null distributions of S, and the median was calculated for each of these distributions, generating a null distribution of S. Median percentile gives the percentile of the null distribution where the observed climate-associated median lies. $p<0.05$ is given in bold for evidence that the climate-associated median S is lower than would be expected by drift alone. Empirical p-value is given for the test that the climate associated median is greater than the non-climate associated distribution of medians. $p<0.05$ is given in bold. $p<0.10$ is underlined.
